# Interactions between Viperin, IRAK1 andTRAF6 couple innate immune signaling to antiviral ribonucleotide synthesis

**DOI:** 10.1101/318840

**Authors:** Arti B. Dumbrepatil, Soumi Ghosh, Ayesha M. Patel, Kelcie A. Zegalia, Paige A. Malec, J. Damon Hoff, Robert T. Kennedy, E. Neil. G Marsh

**Author notes:** Corresponding author: Dr. Neil Marsh, Department of Chemistry, 930 N. University Ave. Ann Arbor MI 48109-1055.

## Abstract

Viperin is a radical S-adenosylmethionine (SAM) enzyme that plays a multifaceted role in the cellular antiviral response. Viperin was recently shown to catalyze the SAM-dependent formation of 3′-deoxy-3′,4′-didehydro-CTP (ddhCTP), which inhibits some viral RNA polymerases. Viperin is also implicated in regulating K63-linked poly-ubiquitination of interleukin-1 receptor-associated kinase-1 (IRAK1) by the E3 ubiquitin ligase TNF Receptor-Associated Factor 6 (TRAF6) as part of the Toll-like receptor-7 and 9 (TLR7/9) innate immune signaling pathways. We show that IRAK1 and TRAF6 activate viperin to efficiently catalyze the radical-mediated dehydration of CTP to ddhCTP. Furthermore, poly-ubiquitination of IRAK1 requires the association of viperin with IRAK1 and TRAF6. Poly-ubiquitination appears dependent on structural changes induced by SAM binding to viperin but does *not* require catalytically active viperin. The synergistic activation of viperin and IRAK1 provides a mechanism that couples innate immune signaling with the production of the antiviral nucleotide ddhCTP.

## Introduction

The innate immune system is involved in the initial detection of pathogens and coordinates the host’s first line of defense against infection (Wu and Chen 2014). Viperin (**V**irus **i**nhibitory **p**rotein, **e**ndoplasmic **r**eticulum-associated, **in**terferon-inducible (also denoted as RSAD2 or Cig5 in humans) is an interferon-inducible protein that is up-regulated in response to viral infection (Fig. 1) (Chin and Cresswell 2001; Helbig and Beard 2014; Seo, Yaneva, and Cresswell 2011). Viperin has been shown to restrict the infectivity of a range of viruses including HIV-1, influenza A, human cytomegalovirus (HCV), Bunyamwera virus, Zika virus and tick-borne encephalitis virus (TBEV) (Helbig et al. 2013; Helbig and Beard 2014; Seo, Yaneva, and Cresswell 2011; Nasr et al. 2012; Helbig et al. 2011; Upadhyay et al. 2014; Panayiotou et al. 2018). An N-terminal membrane-associated domain localizes viperin to the endoplasmic reticulum and lipid droplets, which are the assembly and replication sites for various viruses (Fenwick et al. 2017; Haldar et al. 2012; Shaveta et al. 2010; Hinson and Cresswell 2009b).

**Figure 1:**
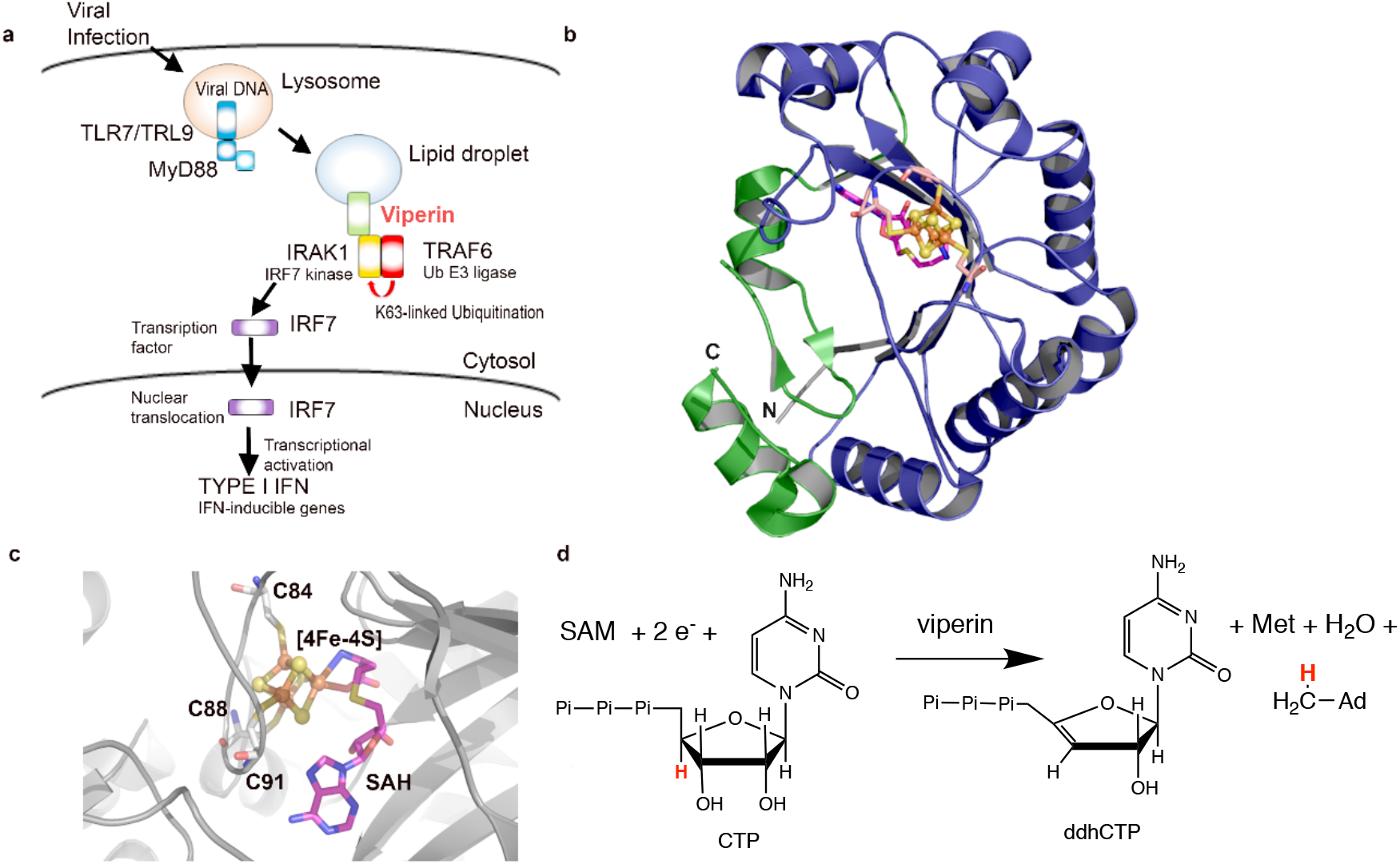
Overview of viperin’s role in TLR7/9 signaling and antiviral ribonucleotide synthesis. **a)** Viperin facilitates the activation of IRAKI for phosphorylation of IRF7 in the TLR7/9 signaling pathway. **b)** Structure of viperin (PDB code 5VSL); the canonical radical SAM domain shown in blue and the viperin-specifìc C-terminal domain in shown green; the [4Fe-4S] cluster is shown as spheres and the structure of SAH as sticks. **c)** Details of the active site showing the [4Fe-4S] cluster (spheres), the ligating cysteine residues of the CxxxCxxC motif and SAH bound (sticks). **d)** SAM-dependent dehydration of CTP to ddhCTP catalyzed by viperin.

Viperin is unusual in being one of only eight radical SAM enzymes found in animals (Landgraf, McCarthy, and Booker 2016). The vast majority of the over 220,000 predicted radical SAM enzymes catalyze reactions unique to microbial metabolism, which are initiated by the reductive cleavage of SAM to generate a 5′-deoxyadenosyl radical (Landgraf et al., 2016; Broderick et al. 2014; Marsh et al., 2004; Frey et al., 2008; Vey and Drennan 2011; Wang et al. 2014). The structure of mouse viperin (Fenwick et al. 2017) shows that it adopts the canonical (βα)6-partial barrel fold observed for radical SAM enzymes (Vey and Drennan 2011), in which the catalytically essential [4Fe-4S] cluster is ligated by the three cysteinyl residues of the hallmark CxxxCxxC motif (Fig. 1b, c) (Fenwick et al. 2017). Very recently it has been shown that viperin catalyzes the formation of a novel antiviral ribonucleotide, 3′-deoxy-3′,4′-didehydro-CTP (ddhCTP; Fig 1d) and that overexpression of viperin in HEK293 causes significant levels of ddhCTP to accumulate (Gizzi et al. 2018). ddhCTP is formed by the dehydration of CTP in a SAM-dependent reaction that is initiated by abstraction of the 4’-hydrogen by the 5’-deoxyadenosyl radical. ddhCTP was shown to be an effective chain-terminating inhibitor of RNA synthesis by viral RNA-dependent RNA polymerases from flaviviruses including dengue virus, west Nile virus and Zika virus, thereby providing an explanation for viperin’s antiviral properties (Gizzi et al. 2018).

The production of anti-viral nucleotides provides one explanation for viperin’s antiviral properties; however, many aspects of the enzyme’s antiviral effects are not readily explained by the production of ddhCTP. For example, it appears that catalytically inactive viperin mutants are still capable of restricting influenza A infection (Vonderstein et al. 2017), and ddhCTP was ineffective as a chain terminator for RNA polymerases from picornaviruses such as poliovirus and human rhinovirus (Gizzi et al. 2018). Furthermore, there is a large body of evidence linking viperin’s antiviral properties to interactions with a variety of host and viral proteins that may play a role in down-regulating or inhibiting metabolic pathways important for viral replication (Wang et al., 2007; Makins et al. 2016; Seo et al. 2011; Vonderstein et al. 2017; Panayiotou et al. 2018; Helbig et al. 2011; Wang et al. 2012). Viperin’s interactions with these proteins is not obviously related to the production of ddhCTP.

In particular, viperin plays an important role in innate immune signaling as a component of the toll-like receptor 7 (TLR7) and TLR9 pathways that leads to type I interferon production (Jiang and Chen 2011; Saitoh et al. 2011). Toll-like receptor 7 (TLR7) and TLR9 are among various pattern recognition receptors that sense viral nucleic acids and induce production of type I interferons by plasmacytoid dendritic cells (pDCs) to protect the host from viral infection (Wu and Chen 2014). As a component of this pathway, viperin stimulates the K63-linked poly-ubiquitination of interleukin-1 receptor-associated kinase (IRAK1) (Wang et al. 2017; Conze et al. 2008; Honda et al. 2004) by the E3 ubiquitin ligase, TRAF6 (Saitoh et al. 2011; Conze et al. 2008). K63-linked poly-ubiquitination, in turn, activates IRAK1 to phosphorylate interferon regulatory factor 7 (IRF7), causing IRF7 to migrate to the nucleus where it activates transcription of type 1 interferons (Honda et al. 2004; Honda et al. 2006) (Fig. 1a).

The fact that viperin is a component of this signaling pathway suggested to us that its enzymatic activity might be regulated by interactions with its partner proteins. To examine this possibility, we reconstituted the interactions between viperin, IRAK1 and TRAF6 that lead to poly-ubiquitination of IRAK1 in HEK 293T cells. Our experiments demonstrate that IRAK1 and TRAF6 activate viperin towards reductive cleavage of SAM and the production of ddhCTP. At the same time, viperin stimulates the poly-ubiquitination of IRAK1 by TRAF6, in a manner that is SAM-dependent. The synergistic activation of these enzymes provides a mechanism to couple innate immune signaling of viral infection to the production of antiviral ribonucleotides.

## Results

It has not proved possible to over-express full-length IRAK1 and TRAF6 in *E. coli*, although various sub-domains of these proteins have been successfully produced (Fu et al. 2018; Ye et al. 2002; Wang et al. 2017). This is most likely because both enzymes are multi-domain human proteins, most of which are notoriously recalcitrant to expression in recombinant bacteria. Partly for this reason, and because it is not clear which domains of these proteins interact with viperin, we decided to study the interactions between viperin, TRAF6 and IRAK1 in a cellular context. Expression in eukaryotic cells allowed viperin to localize to the ER membrane and lipid droplets, which studies suggest may be important for its function (Hinson and Cresswell 2009a; Helbig et al. 2011), and allowed the effect of viperin on the poly-ubiquitination of IRAK1 by TRAF6 to be examined.

Viperin, IRAK1 and TRAF6 were co-transfected in HEK 293T cells and their expression analyzed by immunoblotting. All three proteins were co-expressed, although as evident in Fig 2a, the cellular levels of viperin were lower when IRAK1 and TRAF6 were co-expressed with viperin. The expression levels of TRAF6 or IRAK1 remained similar, regardless of whether the other two enzymes were co-expressed (Fig. 2b, c). Similar patterns of expression were observed when the catalytically inactive mutant, viperinΔ3C, (in which the cysteine ligands that bind the iron-sulfur cluster are mutated to alanine) was co-expressed with IRAK1 and TRA6 (Supplementary Fig 2).

**Figure 2:**
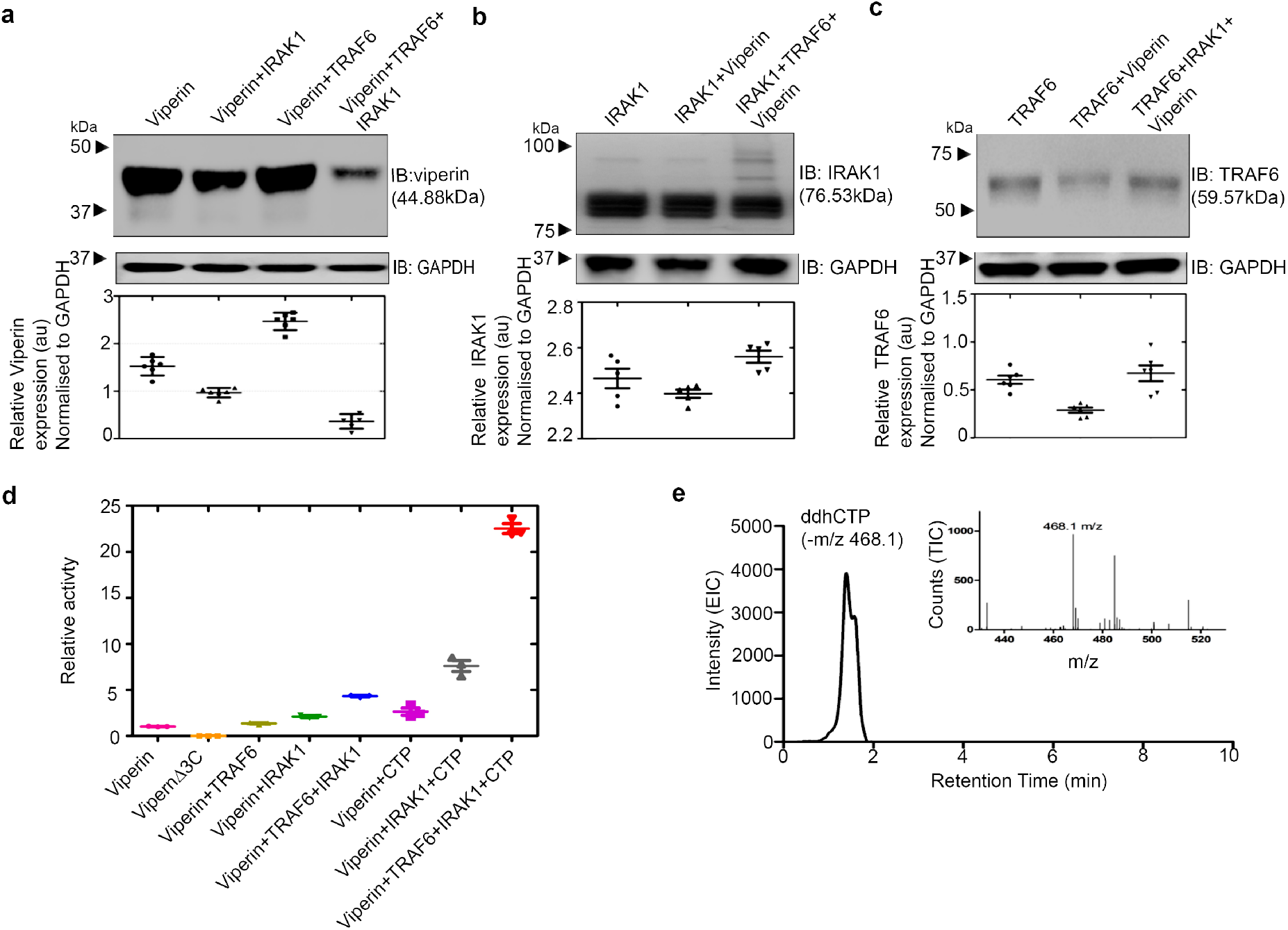
IRAK1 and TRAF6 synergistically activate viperin. HEK293T cells were transfected with the indicated expression constructs. Cells were harvested 30 h post transfection. and lysates from equal numbers of cells were subjected to immunoblotting using the antibodies indicated. GAPDH served as the loading control. **a)** Analysis of viperin expression in the presence or absence of IRAK1 and TRAF6. Co-expression of viperin with IRAK1 and TRAF6 results in a significant decrease in its levels. **b)** Analysis of IRAK1 expression in the presence or absence of viperin and TRAF6; no significant change in IRAK1 expression is seen, however, co-expression with viperin and TRAF6 results in high Mr isoforms of IRAK due to poly-ubiquitination, as discussed in the main text. **c)** Analysis of TRAF6 expression in the presence or absence of IRAK1 and viperin; no significant changes in TRAF6 levels are observed. Quantitation of relative protein expression levels for viperin, IRAK1 and TRAF6 expression relative to GAPDH is presented as mean +/− S.E.M. (n=3) with * indicating p < 0.05, (Student’s t-test for independent samples). d) Quantification of viperin activity in cell extracts. The amount of 5’-dA formed in 1 h, normalized for the amount of viperin present in the cell extracts, is plotted relative to the viperin-only sample = 1.0. The data represent the mean and standard deviation of three independent biological replicates with three technical replicates of each measurement. e) ddhCTP formation in HEK293T cells expressing viperin, IRAK1 and TRAF6 analyzed by LC-MS; *Inset* mass spectrum extracted from chromatograph confirming *m/z* of ddhCTP.

### IRAK1 and TRAF6 synergistically activate viperin

As enzymes in signaling pathways typically function by activating or repressing the activity of other enzymes, the involvement of viperin in the TLR7/9 signaling pathways suggested that the enzymatic activity of viperin may be regulated by IRAK1 and/or TRAF6. Therefore, we examined the effects of IRAK1 and TRAF6 co-expression on the enzymatic activity of viperin. Cell extracts were prepared under anaerobic conditions (due to the *in vitro* oxygen-sensitivity of radical SAM enzymes) from HEK 293T cells co-transfected with viperin, IRAK1 and TRAF6. The enzymatic activity of viperin was quantified by measuring the formation of 5′-dA, as described in the Methods section and the amount of viperin present in the cell extracts was quantified by immunoblotting, (Supplementary Fig. 3)

When assayed in the absence of exogenous CTP, cell extracts expressing only viperin exhibited low levels of activity, with an apparent turnover number, *k*_obs_ = 0.94 ± 0.05 h^-1^ (Fig. 2d, Supplementary Table 1). This basal activity is likely due to low concentrations endogenous CTP in the cell extract. Negligible amounts of 5′-dA were formed in cell extracts lacking viperin or in cells expressing viperinΔ3C. Surprisingly, when the co-substrate CTP (300 μM) was added to the assay, only a modest 2.4-fold increase in activity (*k*_obs_ = 2.38 ± 0.05 h^-1^) was observed. For comparison, previous experiments on N-terminal truncated rat viperin, expressed and purified from *E. coli*, had found CTP stimulated 5′-dA formation ~ 130-fold (Gizzi et al. 2018).

In contrast, when lysates prepared from cells co-expressing IRAK1 and TRAF6 were assayed, viperin activity with CTP as substrate increased ~10-fold (Fig 2d). Under these conditions *k*_obs_ = 21.4 ± 1.6 h^-1^, which is 2-fold higher than the apparent *k*_cat_ reported for truncated rat viperin (Gizzi et al. 2018). In the absence of exogenous CTP, the basal level of viperin activity was increased 4-fold (*k*_obs_ = 4.30 ± 0.21 h^-1^) by co-expression with IRAK1 and TRAF6, consistent with these proteins activating viperin. Expression of viperin with IRAK1 alone resulted in lower levels of activation, whereas co-expression of viperin with TRAF6 alone had no effect on viperin activity. Further experiments confirmed that CTP was converted into ddhCTP by viperin (Fig. 2e) and that when deuterated CTP was used as the substrate, deuterium was transferred to 5′-dA, which is consistent with the postulated mechanism for ddhCTP formation. We also note that in cell extracts ATP, but not GTP or UTP, had a similar stimulatory effect on 5′-dA production (Supplementary Fig. 4, and Supplementary Table 1). However, ATP was not a substrate for N-terminal-truncated viperin expressed and purified from *E. coli;* nor was deuterium from labeled ATP incorporated into 5′-dA (Supplementary Fig. 4). We believe that the stimulatory effect of ATP in cell extracts likely arises from phosphorylation of the endogenous pool of CMP and CDP in the cell extracts, with ATP serving as the phosphate donor.

### IRAK1 mediates interactions between Viperin and TRAF6

To examine whether the activating effect of IRAK1 and TRAF6 results from interactions between viperin, IRAK1 and TRAF6, rather than indirectly through other intermediary protein(s) or metabolites, we conducted immunoprecipitation experiments using FLAG-tagged viperin and Myc-tagged IRAK1 as the bait protein. Immunoprecipitation experiments were conducted on HEK 293T cells transfected with all three proteins and with extracts of cells singly transfected with viperin, IRAK1 or TRAF6. For singly transfected cell extracts these were combined in a 1:1:1 ratio and incubated for 30 min at 4° C before being subjected to immunoprecipitation. In both cases, anti-FLAG M2 affinity gel was then used to immunoprecipitate viperin. In case of IRAK1 as a bait protein, anti-Myc antibodies were used to immunoprecipitate. Both experiments yielded similar results (Fig. 3).

**Figure 3:**
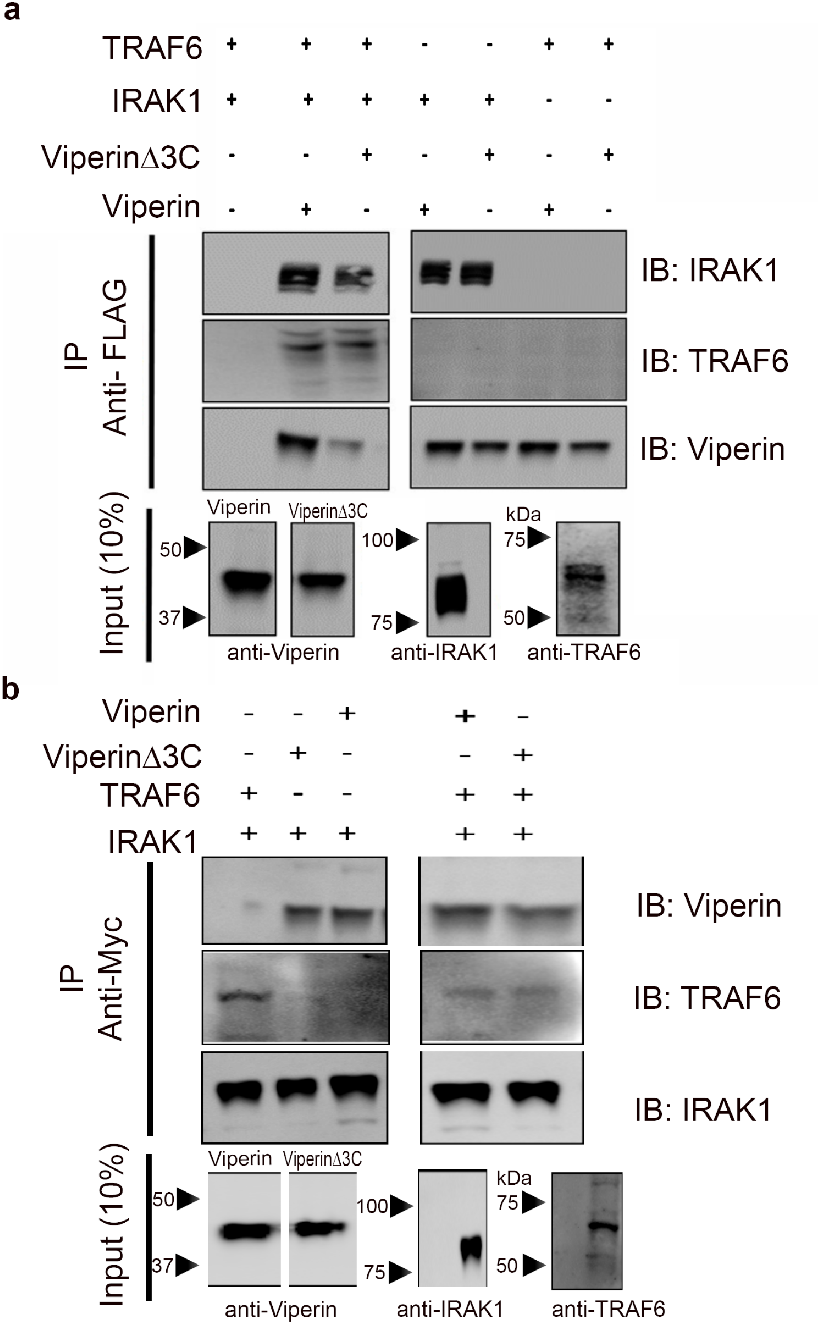
IRAK1 mediates formation of the complex between viperin, IRAK1 and TRAF6. Immuno-tagged genes were transfected into HEK 293T cells and cell extracts were prepared 30 h post transfection. FLAG-viperin or FLAG-viperinΔ3C cell extracts were mixed with IRAK1 and TRAF6 cell extracts in a ratio of 1:1:1. Proteins were immunoprecipitated with either anti-FLAG (viperin) or anti-Myc (IRAK1) antibodies and analyzed by immunoblotting with the indicated antibodies as described in the methods section. **a)** Immunoprecipitation of viperin indicates that viperin binds IRAK1 but not TRAF6. **b)** Immunoprecipitation of IRAK1 indicates that IRAK1 binds both viperin and TRAF6. Mutations in the radical SAM domain (viperinΔ3C) do not affect ability of viperin to interact with IRAK1. Representative images are shown from two independent experiments. Cytosolic extracts were also immunoblotted to confirm expression levels of each individual protein of interest (10% input). For details see Methods.

With viperin as bait, IRAK1 co-precipitated independently of TRAF6; however, TRAF6 only co-precipitated in presence of both viperin and IRAK1 (Fig 3a, Supplementary Fig. 5a). Consistent with these results, when IRAK1 was used as bait both viperin and TRAF6 were co-precipitated independently of each other (Fig. 3b, Supplementary Fig. 5b). These observations imply that IRAK1 binds to both TRAF6 and viperin to mediate complex formation, whereas viperin and TRAF6 do not independently associate with each other. Similar results were obtained when viperinΔ3C was used as the bait protein, which suggests that the iron-sulfur cluster is unnecessary for IRAK1 to bind viperin.

The N-terminal domain of viperin serves to localize the enzyme to the cytoplasmic face of lipid droplets and the endoplasmic reticulum (ER) (Hinson and Cresswell 2009b), whereas TRAF6 and IRAK1 are cytosolic enzymes (Blasius and Beutler 2010). Consistent with the immunoprecipitation results, we observed that co-expression of viperin with TRAF6 and IRAK1 caused these enzymes to re-localize to the ER membrane as determined by immunofluorescence microscopy of fixed and immune-stained cells (Supplementary Fig. 6).

### Poly-ubiquitination of IRAK1 requires viperin

Given that IRAK1 and TRAF6 appeared to activate viperin towards the production of ddhCTP, we were interested to know whether, conversely, viperin stimulated the poly-ubiquitination of IRAK1 by TRAF6. Viperin was initially implicated in the K63-linked poly-ubiquitination of IRAK1 by TRAF6 based on studies that used mouse-derived viperin^+/+^ and viperin^-/-^ pDC cells. The TLR7/9 signaling pathways were stimulated in these cells with either dsRNA or lipopolysaccharides to induce viperin expression (Saitoh et al. 2011; Jiang and Chen 2011). However, these studies did not address the mechanism by which viperin in facilitates immune signaling.

Preliminary evidence that viperin stimulates the poly-ubiquitination of IRAK1, was provided by the observation of high molecular weight isoforms of IRAK1 in immunoblots of cells transfected with viperin, IRAK1 and TRAF6 (Fig. 2b). When immunoblots were probed with anti-ubiquitin antibodies, viperin was found to significantly stimulate poly-ubiquitination of IRAK1 Similar results were found when IRAK1 was first immuno-precipitated from cell extracts and the recovered protein then analyzed by immunoblotting with anti-ubiquitin antibodies, thereby confirming that viperin specifically stimulates IRAK1 poly-ubiquitination, rather than causing a general increase in cellular protein ubiquitination (as discussed below) (Fig. 4, Supplementary Fig. 7).

**Figure 4:**
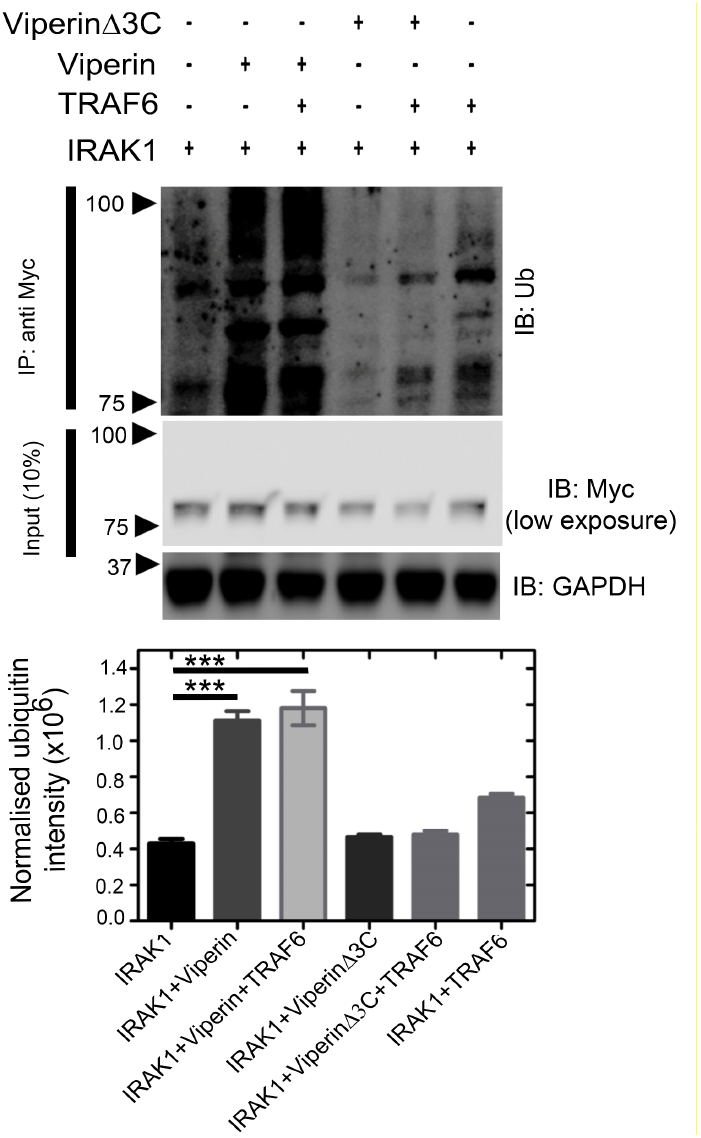
Viperin stimulates poly-ubiquitination of IRAK1. HEK293T cells were transfected with genes encoding IRAK1, TRAF6, viperin or viperinΔ3C as indicated. 30 h post transfection cell lysates were prepared and analyzed by immunoprecipitation (IRAK1 as bait protein) followed by immunoblotting with anti-ubiquitin antibody. GAPDH was used as a loading control. A representative blot is shown. A significant increase in ubiquitinated proteins is only observed in cells expressing viperin and IRAK1; viperinΔ3C fails to stimulate ubiquitination. Ubiquitination was quantified for each condition using ImageJ software and compared. presented as mean +/− S.E.M. (n=3) with *** indicating p < 0.001, (Student’s t-test for independent samples).

Interestingly, co-expression of TRAF6 with viperin did not change the level of IRAK1 ubiquitination significantly (Fig. 4). This suggests that the complex of viperin with IRAK1 either recruits endogenous TRAF6 and/or other E3 ubiquitin ligases to poly-ubiquitinate IRAK1 (Ordureau et al. 2008; Saitoh et al. 2011). Overall, the fraction of over-expressed IRAK1 converted to high molecular weight isoforms remained relatively small. This observation reflects the fact that ubiquitination is a dynamic process in which de-ubiquitination pathways also operate; furthermore, the high, non-physiological concentrations of IRAK1 produced by transfection may exceed the capacity of the cellular ubiquitination machinery. In contrast, the inactive viperinΔ3C mutant failed to stimulate poly-ubiquitination of IRAK1 (Fig. 4).

We note that the activation of IRAK1 by poly-ubiquitination is essential for phosphorylation of IRF7, which is the penultimate step in TLR7/9 signaling (Honda et al. 2004; Honda, Takaoka, and Taniguchi 2006). Interestingly, very little poly-ubiquitination of IRAK1 by TRAF6 was evident in the absence of viperin, even though these proteins were expressed at very high (non-physiological) cellular concentrations (Supplementary Fig. 6). This finding underscores the necessity of the interaction between viperin, IRAK1 and TRAF6 for poly-ubiquitination of IRAK1 to occur.

### Depleting cellular SAM levels reduces the half-life of viperin

To investigate whether SAM was required for viperin to stimulate poly-ubiquitination of IRAK1, we examined the effect of depleting cellular SAM levels using cycloleucine, which is an inhibitor of SAM synthase (Caboche and Hatzfeld 1978). As discussed below, initial observations indicated that cycloleucine does inhibit the poly-ubiquitination of IRAK1. However, when we investigated the effect of cycloleucine (50 mM) on the levels of cellular viperin, we observed that depletion of SAM with cycloleucine significantly reduced the half-life of viperin in the cells (Fig. 5a). This observation suggests that, as is typical of ligands binding to proteins, the binding of SAM to viperin stabilizes its structure, as the half-life of cellular proteins is generally correlated with their stability (Parsell and Sauer 1989; Foit et al. 2009; Sanchez-Ruiz 2010). Therefore, the inhibition of IRAK1 poly-ubiquitination by cycloleucine could be due either to reduced viperin levels, or a requirement for SAM.

**Figure 5:**
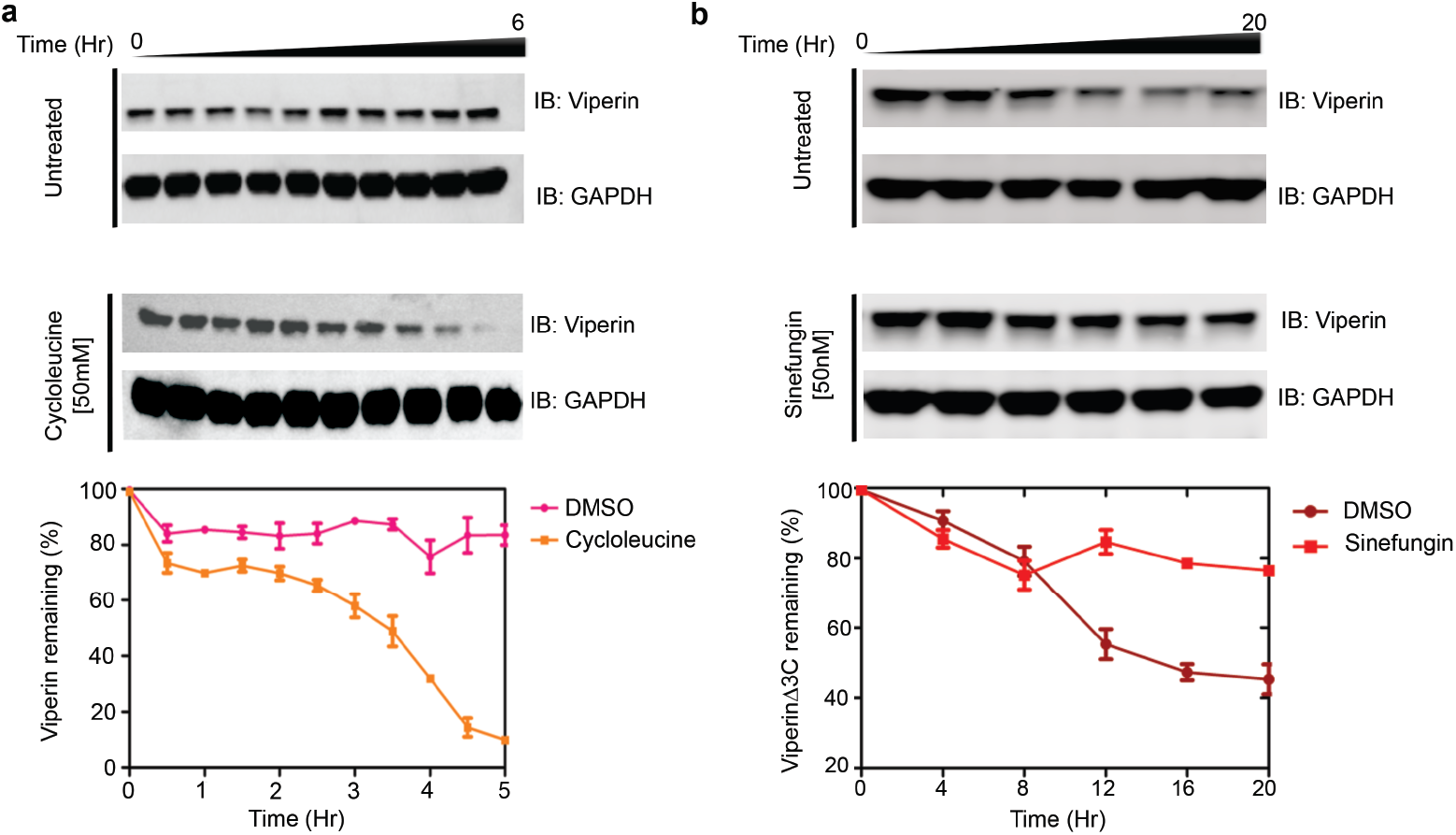
Binding SAM stabilizes viperin *in vivo*. **a)** HEK 293T cells were transfected with viperin and 20 h post transfection 50 mM cycloleucine, a SAM synthase inhibitor, added to the medium. At various time points cells were harvested and viperin expression analyzed by immunoblotting. Depletion of cellular SAM by levels cycloleucine significantly decreases viperin half-life. **b)** HEK 293T cells were transfected with viperinΔ3C and 12 h post transfection 50 nM sinefungin added to the medium. At various time points cells were harvested and viperin expression analyzed by immunoblotting. Sinefungin stabilizes viperinΔ3C compared to DMSO control. *Lower panels* Band intensities were quantified relative GAPDH and expressed as mean ± s.d. from three independent experiments.

We also examined the stability of the viperinΔ3C mutant, which is neither able to cleave SAM nor stimulate IRAK ubiquitination. When expressed in HEK293T cells, we found that viperinΔ3C had a significantly shorter half-life than wild-type viperin, even in the absence of cycloleucine (Fig. 5b). This observation suggested that removing the iron-sulfur cluster significantly destabilizes viperin’s structure, again leading to more rapid degradation.

To investigate the *in vivo* stability of viperinΔ3C further, we examined the effect of including sinefungin in the cell culture medium. Sinefungin is a tight-binding competitive inhibitor of many SAM-dependent enzymes (Borchardt et al. 1979) including radical SAM enzymes (Farrar et al. 2010). We first confirmed that sinefungin binds to viperin by demonstrating that it acts as an effective inhibitor of both purified viperin (recombinantly expressed and purified from *E. coli*) and of viperin in lysates of transfected HEK 293T cells (Supplementary Fig. 8). When HEK 293T cells transfected with viperinΔ3C were cultured in the presence of 50 nM sinefungin, the cellular stability of viperinΔ3C were restored to that of wild-type viperin (Fig. 5b). This observation implies that sinefungin stabilizes the structure of viperinΔ3C in the cell. Sinefungin treatment had no effect on the expression levels of wildtype viperin (Supplementary Fig. 9).

These results suggest that both the iron-sulfur cluster and SAM contribute to the stability of viperin’s structure, thereby protecting it from degradation. The absence of the iron-sulfur cluster in viperinΔ3C presumably weakens or prevents SAM from binding, whereas sinefungin, which would not, in any case, coordinate to the iron-sulfur cluster, binds independently of the iron-sulfur cluster thereby explaining the stabilizing effect of sinefungin on viperinΔ3C.

### Sinefungin restores the ability of viperinΔ3C to stimulate poly-ubiquitination of IRAK1

The ability of sinefungin to stabilize the viperinΔ3C mutant in the cell, suggested that sinefungin might also restore the viperinΔ3C mutant’s ability to stimulate poly-ubiquitination of IRAK1 by TRAF6. To test this hypothesis, HEK293T cells were co-transfected with either viperin or viperinΔ3C, IRAK1 and TRAF6 and 12 h post-transfection treated with either cycloleucine (50 mM) or sinefungin (50 nM, final concentration). The cells were cultured for a further 8 h, harvested and IRAK1 immunopurified from the cell extracts using anti-Myc-tagged magnetic beads. Poly-ubiquitinated forms of IRAK1 were then detected by immunoblotting with anti-ubiquitin antibody (Fig. 6a).

**Figure 6:**
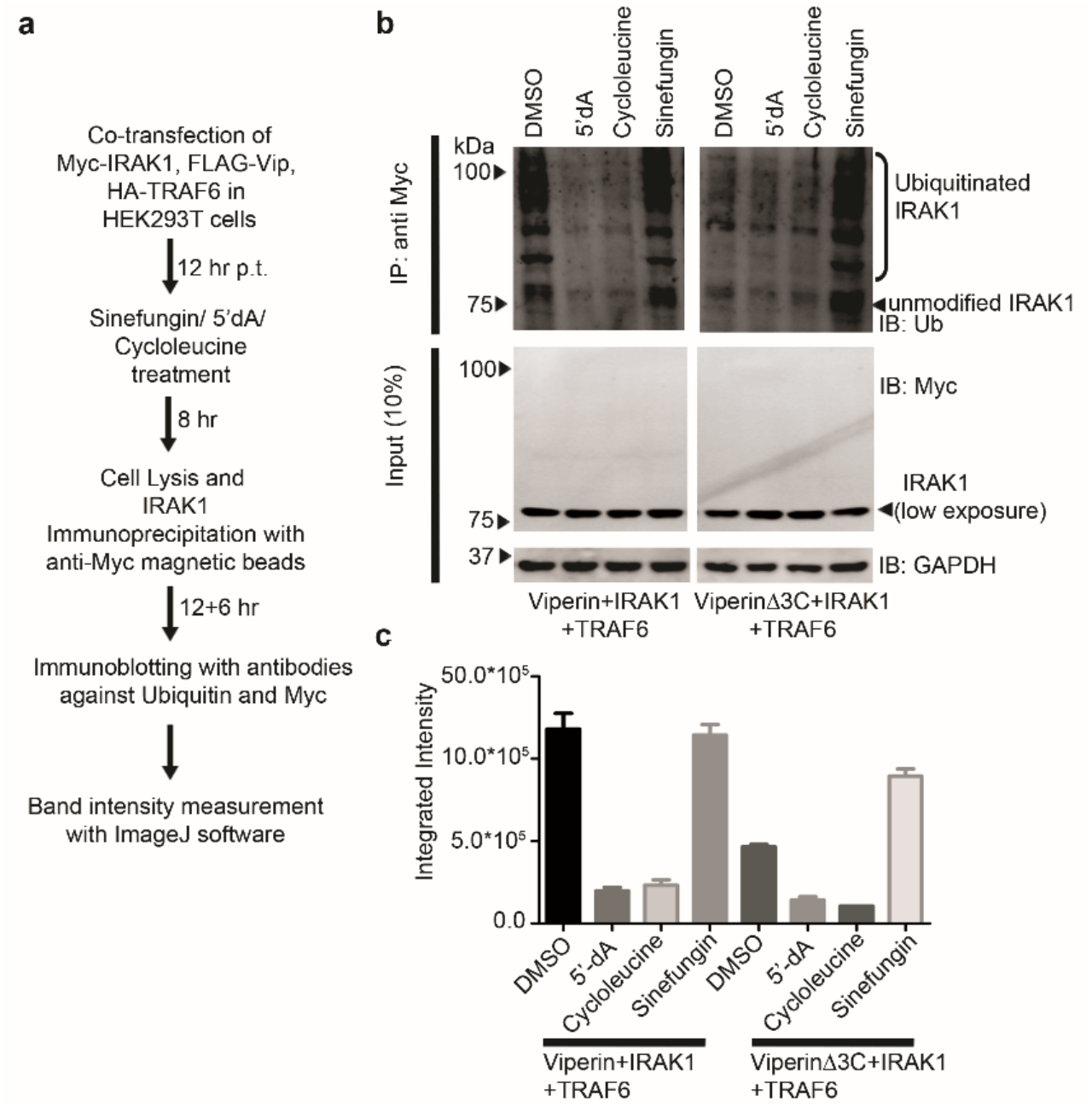
Effects of cycloleucine, 5’-dA and sinefungin on poly-ubiquitination-stimulating activity of viperin. **a)** Flow chart describing experimental protocol. **b)** Cell lysates were immunoprecipitated with the Myc antibody (magnetic beads), followed by immunoblotting with anti-ubiquitin antibodies. Expression of transiently transfected proteins was confirmed by immunoblotting with c-Myc antibody. GAPDH served as a loading control. For wildtype viperin, cycloleucine or 5’-dA inhibit the ubiquitination of IRAK1; in contrast, sinefungin restores the ability of viperinΔ3C, to stimulate the ubiquitination of IRAK1. **c)** Quantification of IRAK ubiquitination levels obtained by integrating intensities of bands for blot using ImageJ software shown in the top panel of b *** indicates statistical significance with *p* < 0.001.

Cycloleucine treatment, as expected, resulted in no poly-ubiquitination of IRAK1 in cells expressing either viperin or viperinΔ3C and IRAK1 and TRAF6 (Fig. 6b). However, sinefungin treatment significantly restored the ability of viperinΔ3C to stimulate poly-ubiquitination of IRAK1 (Fig. 6b). These observations imply that SAM (or a SAM analog) must be bound to viperin for it to form a stimulatory complex with IRAK1 and TRAF6. Importantly, they also demonstrate that the stimulatory effect of viperin on IRAK1 poly-ubiquitination does not depend on the radical-SAM activity of viperin *per se*.

Based on these observations, we hypothesized further that 5’-dA, as the product of SAM cleavage, might inhibit viperin’s ability to stimulate poly-ubiquitination of IRAK1. To test this hypothesis, HEK 293T cells were co-transfected with wild-type viperin, TRAF6 and IRAK1 and treated with 5’-dA (5 μM final concentration). The treated cells were harvested after 12 h and IRAK1 immunopurified from the cell extracts using anti-Myc-tagged beads, and then analyzed by immunoblotting for the presence of poly-ubiquitinated IRAK1. In this case, we observed that 5’-dA significantly reduced the level of poly-ubiquitinated IRAK1 (Fig. 6b), suggesting that 5’-dA inhibits wild-type viperin’s ability to stimulate poly-ubiquitination of IRAK1. Control experiments established that 5’-dA is not a general inhibitor of protein ubiquitination, and that did 5’-dA did not affect the expression of viperin, IRAK1 or TRAF6 (Supplementary Fig. 10).

## Discussion

The recent discovery that viperin catalyzes the synthesis of ddhCTP (Gizzi et al. 2018) has provided an important advance in our understanding of the mechanism by which viperin exerts its antiviral effects. However, although ddhCTP was shown to derive its antiviral effects by acting as an effective chain terminator of RNA-dependent RNA polymerases from flaviviruses such as Zika virus, other RNA polymerases from RNA viruses that viperin is known to restrict were found to be insensitive to ddhCTP (Gizzi et al. 2018). Furthermore, in several cases the radical SAM activity of viperin appears to be dispensable for its antiviral activity (Makins et al. 2016; Vonderstein et al. 2017; Helbig et al. 2013).

A number of studies suggest that interactions between viperin and a variety of host and viral proteins are important for its antiviral effects, although the evidence has tended to rely on indirect measures of viperin’s activity such as the reduction of viral titer or viral RNA levels. By reconstituting the interactions between viperin, IRAK1 and TRAF6 in HEK 293T cells we have been able to examine a specific, biochemically well-defined function of viperin: the activation of IRAK1 by poly-ubiquitination. Our studies show that the enzymatic activity of viperin is increased by ~10-fold by co-expression with IRAK1 and TRAF6, whereas IRAK1 is not efficiently poly-ubiquitinated *in vivo* unless viperin is also co-expressed. Thus, the activities of viperin and IRAK1 appear to be synergistically regulated activities through protein-protein interactions between viperin, IRAK1 and TRAF6.

Given viperin’s role as a component of the TLR7/9 signaling pathways it might be expected that other proteins in the pathway would regulate its enzymatic activity. Regulation may prevent ddhCTP levels accumulating to the point where ddhCTP interferes with cellular RNA polymerases needed to transcribe other proteins involved in the antiviral response. The localization of viperin to the ER-membrane and lipid bodies, which are the sites of flavivirus assembly (Fitzgerald 2011; Helbig et al. 2013; Pena Carcamo et al. 2017), may enhance viperin’s antiviral effect by insuring that ddhCTP is produced at high local concentrations at the site of viral RNA synthesis.

Although our studies have focused on viperin’s interactions with IRAK1 and TRAF6, evidence from other studies (Wang, Hinson, and Cresswell 2007; Makins et al. 2016; Seo et al. 2011; Vonderstein et al. 2017; Panayiotou et al. 2018; Helbig et al. 2011; Wang et al. 2012) indicates that there are many other proteins that interact with viperin and thus might similarly regulate its activity. It is clear that the Fe-S cluster is required for viperin to catalyze the formation of ddhCTP; however, we have shown that the radical-SAM activity is *not* required for viperin to stimulate IRAK1 poly-ubiquitination, because the stimulatory effect of the viperinΔ3C mutant can be rescued by the SAM analog, sinefungin. This dual function of viperin resolves the numerous apparently contradictory observations, described above, regarding whether the radical-SAM activity of viperin is required for its antiviral effects.

The opposing stimulatory and inhibitory effects of sinefungin and 5’-dA, respectively, on IRAK1 poly-ubiquitination suggest that the reductive cleavage of SAM, and concomitant formation of ddhCTP, might serve as a mechanism by which viperin could regulate the ubiquitination of IRAK1 (Fig. 7). In the absence of SAM, viperin adopts a conformation that is unable to stimulate ubiquitination of IRAK1. Upon binding SAM, viperin undergoes a conformational change to an active state that stimulates K63-linked poly-ubiquitination of IRAK1 by TRAF6 (or other E3 ligases), eventually leading to transcription of type I interferons (Saitoh et al. 2011). Once the complex between IRAK1, TRAF6 and viperin is formed, viperin is activated to cleave SAM and at the same time is able to catalyze the formation of ddhCTP and 5’-dA. Completion of the catalytic cycle returns viperin to its inactive state, preventing further poly-ubiquitination of IRAK1.

**Figure 7:**
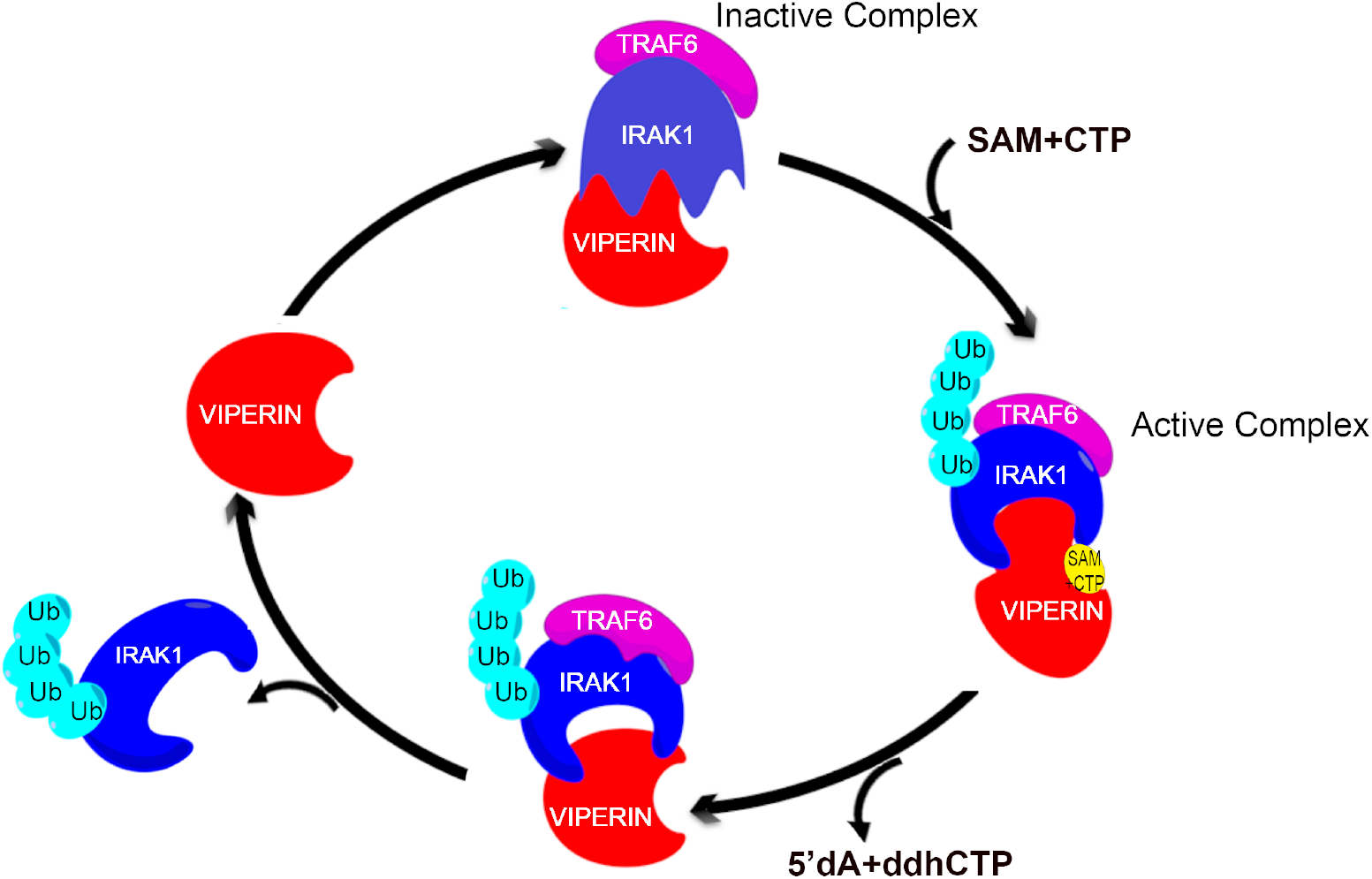
Synergistic activation of viperin, IRAK1 and TRAF6 couples the production of antiviral ribonucleotides with innate immune signaling. *Clockwise from the top:* binding of SAM and CTP to the viperin-IRAK1-TRAF6 complex activates the complex towards poly-ubiquitination of IRAK1. Activated viperin catalyzes the reaction of SAM with CTP to form 5’-dA and ddhCTP and concomitantly switches the complex from the ubiquitination-active state to ubiquitination-inactive state. Poly-ubiquitinated IRAK1 dissociates from the complex, leading to up-regulation of INF-1. For discussion see the text.

Our measurements of viperin activity in cell extracts indicate that viperin converts CTP to ddhCTP with *k*_obs_, = 21.4 ± 1.6 h^-1^, which is about twice as fast as *in vitro* kinetic data reported previously (Gizzi et al. 2018), but still rather slow by comparison with most enzyme reactions. However, the slow kinetics of viperin make this reaction well suited to regulate a signaling pathway in a manner analogous to signal regulation by hetero-trimeric G proteins (Neer 1995; Dohlman and Thorner 1997). We note that this constitutes a new mode of regulating cellular protein activity by radical-SAM enzymes.

In conclusion, our experiments provide evidence for a regulatory mechanism that links the production of the newly identified antiviral nucleotide, ddhCTP, by viperin to transduction of innate immune signaling through K63-linked poly-ubiquitination of IRAK1 by TRAF6. In this manner, the up-regulation of various proteins that constitute the cellular antiviral response is coordinated with the production of a small molecule inhibitor of viral replication. Protein ubiquitination plays multiple roles in cellular proteostasis and signal transduction (Bhoj and Chen 2009; Hu and Sun 2016). We suggest that the wide-ranging roles that viperin plays in the antiviral response may, in part, result from the ability of the enzyme to recruit and activate a subset of the many E3 ubiquitin ligases present in the cell to ubiquitinate target proteins, thereby regulating their cellular levels in response to viral infection.

## Materials and Methods

### Cell lines

The HEK293T cell line was obtained from ATCC.

### Antibodies

Rabbit polyclonal RSAD2/ Viperin antibody (11833-1-AP) was obtained from Protein Tech. Rabbit polyclonal IRAK1 antibody (PA5-17490) was obtained from Thermo Scientific. Rabbit polyclonal Ubiquitin antibody (sc-9133) and mouse monoclonal TRAF6 antibody (sc-8409) were purchased from Santa Cruz. Goat anti-rabbit (170-6515) and anti-mouse (626520) Ig secondary Abs were purchased from BioRad and Life Technologies respectively. Rabbit polyclonal GAPDH (TAB1001) was purchased from Thermo Scientific and Mouse monoclonal GAPDH antibody (6C5) was obtained from EMD Millipore.

### Plasmids

Synthetic genes encoding human viperin, IRAK1 and TRAF6 (GenBank accession numbers AAL50053.1, NM145803, NM001569 respectively) were purchased from GenScript. For details see supplementary information.

### Reagents

The sources of other reagents were as described previously (Makins et al. 2016). Sinefungin (567051-2MG-M) was purchased from Sigma Aldrich. Nucleotide substrates were purchased from: ATP (Adenosine 5′-triphosphate Disodium Salt Trihydrate, Fisher Scientific BP413-25) GTP (Guanosine 5′-triphosphate sodium salt hydrate ≥95%; Sigma G8877), UTP (Uridine 5′-triphosphate, trisodium salt hydrate, 90%; Acros Organics AC226310010) and CTP (Cytidine 5′-triphosphate, disodium salt hydrate, 95%; Acros 0rganics-226225000). Deuterium labelled ATP (Adenosine-2,8-d2,1′,2′,3′,4′,5′,5′-d6 5′-triphosphate sodium salt, 738034-1MG) and CTP (Cytidine-5,6-d2,1′,2′,3′,4′,5′,5′-d6 5′-triphosphate sodium salt, 738042-1MG) were purchased from Sigma Aldrich. Pierce™ Anti-c-Myc Magnetic Beads (88842) were purchased from ThermoFisher Scientific.

### Transfection

HEK 293T cells, cultured as described above, were transiently transfected using FuGENE^®^ HD (Promega) following the manufacturer’s instructions.

### Immunofluorescence analyses

Cells were grown on poly-L-lysine-coated coverslips to 30–40% confluence, transfected with plasmids expressing Viperin, IRAK1 and/or TRAF6 and incubated for 30 h. Cells were fixed with 4% paraformaldehyde, permeabilized with 0.05% Triton X-100 dissolved in PBS, and washed three times with PBS containing 0.1% Tween20. The fixed cells were stained with the appropriate antibodies. Primary antibodies were diluted in PBS containing, 1% FBS and 0.1% Tween 20. Viperin was detected by using mouse monoclonal anti-viperin (Abcam) diluted 1:250, and calnexin was detected with rabbit polyclonal antibody (Abcam) diluted 1:250. IRAK1 was detected by using rabbit polyclonal IRAK1 antibody (Thermo Scientific) diluted 1:250 and mouse monoclonal c-Myc Tag antibody (Thermo Scientific) diluted 1:100. TRAF6 was detected by using mouse monoclonal TRAF6 antibody (Santa Cruz) diluted 1:100. After incubation at room temperature for 1 h or overnight at 4°C, the coverslips were washed with PBS containing 0.1% Tween20 and treated with Alexa Fluor 647-conjugated goat anti-mouse (Life Technologies) and Alexa Fluor 488-conjugated goat anti-rabbit (Abcam) secondary antibodies at a dilution of 1:400 at room temperature for 2 h. The coverslips were washed three times with PBS containing 0.1% Tween20 and mounted in ProLong™ Gold Antifade Mountant (Molecular Probes). Images were acquired with an Olympus IX81 microscope with a 60× 1.49NA objective on an Andor iXON Ultra EMCCD camera. 488 nm (Coherent Cube 488-50) and 640 nm (Coherent Cube 640–100) laser excitation was aligned in HILO imaging mode for axial sectioning using an Olympus cell^TIRF module.

### Immunoblotting

Cells were lysed in TBST buffer (20 mM Tris (pH 7.5), 500 mM NaCl, 0.05% Tween-20) containing protease inhibitors (SIGMAFAST™ Protease Inhibitor Tablets, S8830; Sigma). Supernatants of lysates were collected and mixed with reducing sample buffer. The supernatants were separated on 10% SDS-PAGE gels. The bicinchoninic acid (BCA) assay (Thermo Scientific) was used to determine the total amount of protein in the lysates. Immunoblotting was performed as describes previously (Makins et al. 2016). Primary antibodies used were directed against viperin (rabbit polyclonal diluted 1:2500), GAPDH (rabbit polyclonal diluted 1:5000), IRAK1 (rabbit polyclonal diluted 1:4000), TRAF6 (mouse monoclonal diluted 1:500) and GAPDH (mouse monoclonal diluted 1:5000). Rabbit polyclonal ubiquitin antibody was used at a dilution of 1:1000. Blots were visualized, and band intensities quantified using a Bio-Rad ChemiDoc Touch imaging system. Integrated density measurements were done using ImageJ software. Quantitative measurements of protein expression levels reported here represent the average of at least three independent biological replicates

### Immunoprecipitation Assays

Cells were rinsed twice with ice-cold PBS, harvested in P40 lysis buffer (50 mM Tris, pH 7.4,1% Triton X-100, 150 mM NaCl, 10% glycerol, 1 mM EDTA, 10 mM NaF, and 0.2 mM phenylmethylsulfonyl fluoride with protease inhibitor cocktail from Sigma), and briefly sonicated. Lysates were collected by centrifugation at 12,000 × *g* for 15 minutes at 4°C. For immunoprecipitation, ratio of suspension to packed gel volume was 2:1. Resin pre-equilibration was done as per the manufacturer’s protocol. Three hundred microliters of a 1:1:1 ratio of cell lysates was added and incubated for 16 hours at 4°C with gentle rotation. Beads were pelleted by centrifugation at 5,000 × *g* for 30 seconds at 4°C and washed three times with washing buffer (50 mM Tris, pH 7.4,150 mM NaCl, 10% glycerol, 1 mM EDTA, 10 mM NaF, and 0.2 mM phenylmethylsulfonyl fluoride with protease inhibitor cocktail from Sigma). Immunocomplexes were eluted by boiling in SDS-PAGE sample buffer, separated by SDS-PAGE, and transferred to a nitrocellulose membrane. Immunoprecipitation using Protein A beads or anti Myc-tagged magnetic beads was performed as per the manufacturers protocol. Membranes were blocked for 2 h at room temperature in TBST buffer (20 mM Tris, pH 7.5, 137 mM NaCl, and 0.1% Tween 20) containing 5% nonfat dry milk, followed by overnight incubation at 4°C in TBST buffer containing 3% nonfat dry milk and the appropriate primary antibody. Membranes were washed three times in TBST and then incubated for 2 h at room temperature with the secondary IgG-coupled horseradish peroxidase antibody. The membranes were washed three times with TBST, and the signals were visualized with enhanced chemiluminescence reagent as described in immunoblotting.

### Statistical analyses

Results from all studies were compared with unpaired two-tailed Student’s t test using GraphPad Prism 5 software. P values less than 0.05 were considered significant.

### Assay of viperin in HEK 293T cell lysates

HEK 293T cells transfected with viperin, and/or IRAK1 and TRAF6 were harvested from one 10 cm diameter tissue culture plate each, resuspended in 500 μl of anoxic Tris-buffered saline (50 mM Tris-Cl, pH 7.6, 150 mM NaCl) containing 1% Triton X-100, sonicated within an anaerobic glovebox (Coy Chamber), and centrifuged at 14,000 g for 10 min. Dithiothreitol (DTT; 5 mM) and dithionite (5 mM) were added to the cell lysate together with CTP (300 μM). The assay mixture was incubated at room temperature for 30 min prior to starting the reaction by the addition of SAM (200 μM). The assay was incubated for 60 min at room temperature, after which the reaction stopped by heating at 95 °C for 10 min. The solution was chilled to 4 °C, and the precipitated proteins were removed by centrifugation at 14,000 rpm for 25 min. The supernatant was then extracted with acetonitrile. Samples were analyzed in triplicate by UPLC-tandem mass spectrometry as described previously (Makins et al. 2016). For details of standard curve construction and calculations refer to Supplementary Information.

## Materials & Correspondence

Correspondence and material requests should be addressed to.

## Author Contributions

A.B.D., S.G., P.A.M., K.Z. and J.D.H. performed the experiments. A.B.D., P.A.M., K.Z. and J.D.H. analyzed the results. S.G. provided reagents. A.B.D., J.D.H., R.T.K. and E.N.G.M. designed the experiments. A.B.D. and E.N.G.M. wrote the paper.

## Acknowledgements

This work was supported in part by NIH grants GM 093088 to E.N.G.M. and DK 046960 to R.T.K.

