## Supplemental figures and methods for "Interactions between Viperin, IRAK1 andTRAF6 couple innate immune signaling to antiviral ribonucleotide synthesis"

### Viperin interacts with IRAK1 and TRAF6 to couple immune signaling to antiviral nucleotide synthesis

[*]Corresponding author:

#### Materials and Methods

**Materials**

Sinefungin (567051-2MG-M) was purchased from Sigma Aldrich. Nucleotide substrates were purchased from: ATP (Adenosine 5'-triphosphate Disodium Salt Trihydrate, Fisher Scientific BP413-25) GTP (Guanosine 5′-triphosphate sodium salt hydrate ≥95%; Sigma G8877), UTP (Uridine 5'-triphosphate, trisodium salt hydrate, 90%; Acros Organics AC226310010) and CTP (Cytidine 5'-triphosphate, disodium salt hydrate, 95%; Acros Organics-226225000). Deuterium labelled ATP (ATP-$D_8$, Adenosine-2,8-d2,1′,2′,3′,4′,5′,5′-d6 5′-triphosphate sodium salt, 738034-1MG) and CTP (CTP-$D_8$, Cytidine-5,6-d2,1′,2′,3′,4′,5′,5′-d6 5′-triphosphate sodium salt, 738042-1MG) were purchased from Sigma Aldrich. The sources of other reagents were as described previously (Makins et al. 2016).

**Plasmids**

The genes encoding human viperin (GenBankTM accession number AAL50053.1) were purchased from GenScript. genes were subsequently amplified via PCR and subcloned into the pcDNA3.1(+) vector (Invitrogen). The ViperinΔ3C mutant was constructed by stepwise mutations of the cysteine residues to alanine using the QuickChange Site-Directed Mutagenensis kit (Agilent). A gene, codon-optimized for expression in *E. coli*, encoding viperin lacking the first 50 amino acids of the N-terminal amphipathic α-helix was purchased from GenScript. The gene construct was housed in a pET28a expression vector and included a N-terminal His-6 tag to facilitate purification (Makins et al. 2016). The plasmids for IRAK1 expression and TRAF6 expression were purchased from GenScript. IRAK1 (NM_001569) was cloned between *EcoR*I and *Xho*I with an N-terminal Myc tag. TRAF6 (NM_145803) was cloned between *EcoR*I and *Xho*I with an N-terminal HA tag (this paper).

**Gene sequence for IRAK1**

5’GAATTCATGGCCGGGGGGCCGGGCCCGGGGGAGCCCGCAGCCCCCGGCGCCCAGCACTTCTT
GTACGAGGTGCCGCCCTGGGTCATGTGCCGCTTCTACAAAGTGATGGACGCCCTGGAGCCCGCC
GACTGGTGCCAGTTCGCCGCCCTGATCGTGCGCGACCAGACCGAGCTGCGGCTGTGCGAGCGCT
CCGGGCAGCGCACGGCCAGCGTCCTGTGGCCCTGGATCAACCGCAACGCCCGTGTGGCCGACCT
CGTGCACATCCTCACGCACCTGCAGCTGCTCCGTGCGCGGGACATCATCACAGCCTGGCACCCT
CCCGCCCCGCTTCCGTCCCCAGGCACCACTGCCCCGAGGCCCAGCAGCATCCCTGCACCCGCCG
AGGCCGAGGCCTGGAGCCCCCGGAAGTTGCCATCCTCAGCCTCCACCTTCCTCTCCCCAGCTTT
TCCAGGCTCCCAGACCCATTCAGGGCCTGAGCTCGGCCTGGTCCCAAGCCCTGCTTCCCTGTGG
CCTCCACCGCCATCTCCAGCCCCTTCTTCTACCAAGCCAGGCCCAGAGAGCTCAGTGTCCCTCC
TGCAGGGAGCCCGCCCCTTTCCGTTTTGCTGGCCCCTCTGTGAGATTTCCCGGGGCACCCACAA
CTTCTCGGAGGAGCTCAAGATCGGGGAGGGTGGCTTTGGGTGCGTGTACCGGGCGGTGATGAGG
AACACGGTGTATGCTGTGAAGAGGCTGAAGGAGAACGCTGACCTGGAGTGGACTGCAGTGAAGC
AGAGCTTCCTGACCGAGGTGGAGCAGCTGTCCAGGTTTCGTCACCCAAACATTGTGGACTTTGC
TGGCTACTGTGCTCAGAACGGCTTCTACTGCCTGGTGTACGGCTTCCTGCCCAACGGCTCCCTG
GAGGACCGTCTCCACTGCCAGACCCAGGCCTGCCCACCTCTCTCCTGGCCTCAGCGACTGGACA
TCCTTCTGGGTACAGCCCGGGCAATTCAGTTTCTACATCAGGACAGCCCCAGCCTCATCCATGG
AGACATCAAGAGTTCCAACGTCCTTCTGGATGAGAGGCTGACACCCAAGCTGGGAGACTTTGGC
CTGGCCCGGTTCAGCCGCTTTGCCGGGTCCAGCCCCAGCCAGAGCAGCATGGTGGCCCGGACAC
AGACAGTGCGGGGCACCCTGGCCTACCTGCCCGAGGAGTACATCAAGACGGGAAGGCTGGCTGT
GGACACGGACACCTTCAGCTTTGGGGTGGTAGTGCTAGAGACCTTGGCTGGTCAGAGGGCTGTG
AAGACGCACGGTGCCAGGACCAAGTATCTGAAAGACCTGGTGGAAGAGGAGGCTGAGGAGGCTG
GAGTGGCTTTGAGAAGCACCCAGAGCACACTGCAAGCAGGTCTGGCTGCAGATGCCTGGGCTGC
TCCCATCGCCATGCAGATCTACAAGAAGCACCTGGACCCCAGGCCCGGGCCCTGCCCACCTGAG
CTGGGCCTGGGCCTGGGCCAGCTGGCCTGCTGCTGCCTGCACCGCCGGGCCAAAAGGAGGCCTC
CTATGACCCAGGTGTACGAGAGGCTAGAGAAGCTGCAGGCAGTGGTGGCGGGGGTGCCCGGGCA
TTCGGAGGCCGCCAGCTGCATCCCCCCTTCCCCGCAGGAGAACTCCTACGTGTCCAGCACTGGC
AGAGCCCACAGTGGGGCTGCTCCATGGCAGCCCCTGGCAGCGCCATCAGGAGCCAGTGCCCAGG
CAGCAGAGCAGCTGCAGAGAGGCCCCAACCAGCCCGTGGAGAGTGACGAGAGCCTAGGCGGCCT
CTCTGCTGCCCTGCGCTCCTGGCACTTGACTCCAAGCTGCCCTCTGGACCCAGCACCCCTCAGG
GAGGCCGGCTGTCCTCAGGGGGACACGGCAGGAGAATCGAGCTGGGGGAGTGGCCCAGGATCCC
GGCCCACAGCCGTGGAAGGACTGGCCCTTGGCAGCTCTGCATCATCGTCGTCAGAGCCACCGCA
GATTATCATCAACCCTGCCCGACAGAAGATGGTCCAGAAGCTGGCCCTGTACGAGGATGGGGCC
CTGGACAGCCTGCAGCTGCTGTCGTCCAGCTCCCTCCCAGGCTTGGGCCTGGAACAGGACAGGC
AGGGGCCCGAAGAAAGTGATGAATTTCAGAGCTGACTCGAG3’

#### Gene sequence for TRAF6

5′GAATTCATGAGTCTGCTAAACTGTGAAAACAGCTGTGGATCCAGCCAGTCTGAAAGTGACTG
CTGTGTGGCCATGGCCAGCTCCTGTAGCGCTGTAACAAAAGATGATAGTGTGGGTGGAACTGCC
AGCACGGGGAACCTCTCCAGCTCATTTATGGAGGAGATCCAGGGATATGATGTAGAGTTTGACC
CACCCCTGGAAAGCAAGTATGAATGCCCCATCTGCTTGATGGCATTACGAGAAGCAGTGCAAAC
GCCATGCGGCCATAGGTTCTGCAAAGCCTGCATCATAAAATCAATAAGGGATGCAGGTCACAAA
TGTCCAGTTGACAATGAAATACTGCTGGAAAATCAACTATTTCCAGACAATTTTGCAAAACGTG
AGATTCTTTCTCTGATGGTGAAATGTCCAAATGAAGGTTGTTTGCACAAGATGGAACTGAGACA
TCTTGAGGATCATCAAGCACATTGTGAGTTTGCTCTTATGGATTGTCCCCAATGCCAGCGTCCC
TTCCAAAAATTCCATATTAATATTCACATTCTGAAGGATTGTCCAAGGAGACAGGTTTCTTGTG
ACAACTGTGCTGCATCAATGGCATTTGAAGATAAAGAGATCCATGACCAGAACTGTCCTTTGGC
AAATGTCATCTGTGAATACTGCAATACTATACTCATCAGAGAACAGATGCCTAATCATTATGAT
CTAGACTGCCCTACAGCCCCAATTCCATGCACATTCAGTACTTTTGGTTGCCATGAAAAGATGC
AGAGGAATCACTTGGCACGCCACCTACAAGAGAACACCCAGTCACACATGAGAATGTTGGCCCA
GGCTGTTCATAGTTTGAGCGTTATACCCGACTCTGGGTATATCTCAGAGGTCCGGAATTTCCAG
GAAACTATTCACCAGTTAGAGGGTCGCCTTGTAAGACAAGACCATCAAATCCGGGAGCTGACTG
CTAAAATGGAAACTCAGAGTATGTATGTAAGTGAGCTCAAACGAACCATTCGAACCCTTGAGGA
CAAAGTTGCTGAAATCGAAGCACAGCAGTGCAATGGAATTTATATTTGGAAGATTGGCAACTTT
GGAATGCATTTGAAATGTCAAGAAGAGGAGAAACCTGTTGTGATTCATAGCCCTGGATTCTACA
CTGGCAAACCCGGGTACAAACTGTGCATGCGCTTGCACCTTCAGTTACCGACTGCTCAGCGCTG
TGCAAACTATATATCCCTTTTTGTCCACACAATGCAAGGAGAATATGACAGCCACCTCCCTTGG
CCCTTCCAGGGTACAATACGCCTTACAATTCTTGATCAGTCTGAAGCACCTGTAAGGCAAAACC
ACGAAGAGATAATGGATGCCAAACCAGAGCTGCTTGCTTTCCAGCGACCCACAATCCCACGGAA
CCCAAAAGGTTTTGGCTATGTAACTTTTATGCATCTGGAAGCCCTAAGACAAAGAACTTTCATT
AAGGATGACACATTATTAGTGCGCTGTGAGGTCTCCACCCGCTTTGACATGGGTAGCCTTCGGA
GGGAGGGTTTTCAGCCACGAAGTACTGATGCAGGGGTATAGCTCGAG3′

#### Supplementary Methods

##### Expression and Purification of recombinant truncated viperin (ΔN-Viperin)

For *in vitro* assays of viperin, a truncated version of the enzyme that lacked the N-terminal amphipathic helix was used. This enzyme, ΔN-viperin was over-expressed an purified from *E. coli* as described previously (Makins et al. 2016).

##### SAM cleavage assays using purified recombinant ΔN-viperin

Enzyme assay reactions were performed under anaerobic conditions in10 mM Tris-Cl buffer pH 8.0, 300 mM NaCl, 10% glycerol and 5 mM sodium dithionite. Assays contained in a total volume of 100 µL: 20 µM protein, 200 µM SAM and either 300 µM CTP, CTP-$D_8$, ATP or ATP-$D_8$. Assays to examine the specificity of hydrogen atom abstraction were performed under similar conditions but with 100 µM ΔN-viperin, 1 mM SAM and 1 mM of CTP or CTP-$D_8$, ATP or ATP-$D_8$ , 5mM dithiothreitol and 5mM sodium dithionite. The assay was incubated for 60 min at room temperature, after which time the reaction stopped by heating at 95 °C for 10 min. The solution was chilled to 4 °C, and the precipitated proteins were removed by centrifugation at 14,000 rpm for 25 min. The supernatant was then extracted with acetonitrile. Samples were analyzed in triplicate by UPLC-tandem mass spectrometry as described in *general methods for mass spectroscopy* section (Supplementary Fig. 3).

##### Inhibition of viperin by sinefungin

To examine the effect of sinefungin on viperin activity, enzyme assay reactions with purified, recombinant ΔN-viperin were performed under the conditions described above. In this case, assays contained in a total volume of 100 µL: 20 µM ΔN-viperin (recombinant purified protein), 200 µM SAM, 300 µM CTP and 800µM sinefungin (SF).

To determine the effect of sinefungin on viperin activity in HEK293T cell lysates, lysates were prepared as described in the main methods section. To 500 $\mu$l of lysate prepared in 50 mM Tris-Cl, pH 7.6, 150 mM NaCl containing 1% Triton X-100 was added 5 mM dithiothreitol and 5 mM sodium dithionite 300 µM CTP and 1 mM sinefungin (final concentrations). The assay mixture was incubated at room temperature for 30 min prior to starting the reaction by the addition of 200 $\mu$M SAM. In both the cases, assay was incubated for 60 min at room temperature, after which time the reaction stopped by heating at 95 °C for 10 min. The solution was chilled to 4 °C, and the precipitated proteins were removed by centrifugation at 14,000 rpm for 25 min. The supernatant was then extracted with acetonitrile. Samples were analyzed in triplicate by UPLC-tandem mass spectrometry as described in *general methods for mass spectroscopy* section. (Supplementary Fig. 9).

**General methods for mass spectroscopy**

20 $\mu$L aliquots of acetonitrile-extracted nucleotides were derivatized with benzoyl chloride as described previously (Wong et al., 2016). Briefly, 10 $\mu$L each of 100 mM sodium carbonate, 2% (v/v) benzoyl chloride, and the internal standard solution were added sequentially, with mixing between each addition. The internal standard solution consisted of $^{13}C_6$-benzoyl chloride derivatized adenosine (Bz-Ado), 5'-deoxyadenosine (Bz-5'dAdoH), and SAM (Bz-SAM) in 20% (v/v) acetonitrile with 1% (v/v) sulfuric acid. Calibration standards were prepared in water and diluted with acetonitrile to match the sample composition.

Samples were analyzed in triplicate by UPLC-tandem mass spectrometry using an Acquity HSS T3 C18 chromatography column (1 mm x 100 mm, 1.8 um, 100 Å pore size) on a Waters nanoAcquity UPLC interfaced to an Agilent 6410B triple quadrupole mass spectrometer. The injection volume was 5 $\mu$L. Mobile phase A was 10 mM ammonium formate with 0.15% formic acid. Mobile phase B was acetonitrile. The flow rate was 100 $\mu$L/min, and the gradient was as follows: initial, 0% B; 0.01 min, 17% B; 0.5 min, 40% B; 2.99 min, 60% B; 3.00 min, 100% B; 3.99 min, 100% B; 4.00 min, 0% B; 5.00 min, 0% B. Positive electrospray ionization mode was used, with the capillary set at 4 kV. The nebulizer pressure was 15 psi, drying gas was 350 °C, and the gas flow was 15 L/min. Detection was performed in dynamic MRM mode, and MRM conditions are listed below (Method 1). Peaks were integrated using Agilent MassHunter Workstation Quantitative Analysis for QQQ, version B.05.00. All peaks were inspected to ensure proper integration.

A modified method was used in the analysis of samples to detect deuterated 5'-deoxyadenosine (Bz-5'dAdoD) using a Acquity HSS T3 C18 chromatography column (2 mm x 100 mm, 1.8 um, 100 Å pore size) on an Agilent 1260 Infinity HPLC interfaced to an Agilent 6410B triple quadrupole mass spectrometer. The injection volume was 5 $\mu$L. Mobile phase A was 10 mM ammonium formate with 0.15% formic acid. Mobile phase B was acetonitrile. The flow rate was 100 $\mu$L/min, and the gradient was as follows: initial, 0 min, 17% B; 1 min, 17% B; 11 min, 100% B; 13 min, 100% B; 13.1 min, 17% B; 18 min, 17% B. Positive electrospray ionization mode was used with the capillary set at 4 kV, nebulizer pressure at 15 psi, drying gas at 350 °C, and the gas flow at 11 L/min. Detection was performed in dynamic MRM mode, and MRM conditions are listed below (Method 2). Peaks were integrated using Agilent MassHunter Workstation Quantitative Analysis for QQQ, version B.05.00. All peaks were inspected to ensure proper integration. For quantitation, deuterated 5'-deoxyadenosine was normalized to the natural isotope abundance found in the calibration standards.

**Summary of MRM methods used to analyze 5’-dA**

| Analyte | Precursor (m/z) | Product (m/z) | Fragmentor voltage (V) | Collision Energy (V) | Cell Accelerator voltage (V) | Retention Time Method 1 (min) | Retention Time Method 2 (min) |
|---|---|---|---|---|---|---|---|
| Bz-SAM | 711.5 | 206 | 140 | 30 | 4 | 2.7 | 11.5 |
| 13C-Bz-SAM | 729.5 | 212 | 140 | 30 | 4 | 2.7 | 11.5 |
| Bz-Ado | 476 | 136 | 120 | 30 | 4 | 3.5 | 13.0 |
| 13C-Bz-Ado | 488 | 136 | 120 | 30 | 4 | 3.5 | 13.0 |
| Bz-5'-dAdoH | 460 | 105 | 120 | 30 | 4 | 4.2 | 14.1 |
| Bz-5’dAdoD | 461 | 105 | 120 | 30 | 4 | -- | 14.1 |
| 13C-Bz-5'-dAdo | 472 | 111 | 120 | 30 | 4 | 4.2 | 14.1 |

**Detection of ddhCTP**

HEK 293T cells transfected with various combinations of viperin, IRAK1 and TRAF6, depending on the experiment, were harvested from one 10 cm diameter tissue culture plate and resuspended in 300 $\mu$l of anoxic Tris-buffered saline (50 mM Tris-Cl, pH 7.6, 150 mM NaCl) containing 1% Triton X-100. The suspension was sonicated within an anaerobic glovebox (Coy Chamber), and centrifuged at 14,000 g for 10 min. Dithiothreitol, 5 mM, and sodium dithionite, 5 mM, were added to the cell lysate together with deuterium-labelled CTP, 1mM. The assay mixture was incubated at room temperature for 30 min prior to starting the reaction by the addition of SAM, 500 $\mu$M. After 60 min the reaction stopped by heating at 95 °C for 10 min. The solution was chilled to 4 °C, and the precipitated proteins were removed by centrifugation at 14,000 rpm for 25 min. The supernatant was then extracted with acetonitrile and concentrated using a SpeedVac concentrator. The extracted material was then redissolved in 30 µL of acetonitrile. 20 µL was directly injected onto C18 ZORSAX ECLIPSE PLVSC18 (2.1x50mm) column on a 6520 Accurate Mass Q-TOF LC/MS. The column was equilibrated with 99.2% mobile phase A (water + 0.1% formic acid) and

0.8% Mobile phase B (acetonitrile + 0.1% formic acid). A gradient of 5% B to 95% B in 16 min was applied with a flow rate of 0.4 ml/min. Detection of CTP and ddhCTP was performed in negative ion mode (ESI-). The mass to charge ratio (m/z) of unlabeled CTP is 482.1 and increases by one mass unit for each deuterium substitution. The m/z of the resulting ddhCTP is 464.1, which increases by one mass unit for each deuterium substitution (Supplementary Fig. 3).

**Standard curve construction for quantifying viperin from cell lysates**

The protein concentration of viperin in lysates of HEK293T cells was determined by quantitative western blot analysis. Cell cultures of the transfected HEK293T cells, were harvested when confluent by resuspending in ice-cold PBS. Cells pellets were washed once in 5 mL PBS and pelleted by centrifugation at 1500 rpm for 5 min at 4°C. Cells were lysed using 500 $\mu$l of SDS lysis buffer (2% SDS in PBS containing a cocktail of protease inhibitors). The lysate was incubated on ice for 10 mins, centrifuged at 543,000 *g* for 1 h at 4 °C to remove DNA and insoluble material and supernatant whole cell lysate was stored at -70°C. The bicinchoninic ccid (BCA) Assay was used to determine the total amount of protein in the lysates. Aliquots of lysates containing known amounts of total protein were subjected to SDS-PAGE and immunoblotted and probed with with anti-viperin antibody. In parallel, known amounts of purified recombinant ΔN-viperin were similarly subjected to SDS-PAGE immunoblotted and probed with with anti-viperin antibody construct a standard curve. This allowed the amount of viperin in the cell extracts to be calculated (Supplementary Fig. 4).

**Supplementary Table 1:**

**Specific activity of viperin in lysates of HEK 293T cells under various conditions; activity assayed by the formation of 5'-dA.**

| Condition | Viperin (pmol) | 5'-dA (pmol) | Turnovers ($h^{-1}$) |
|---|---|---|---|
| Viperin | 1.38 ± 0.07 | 1.30 ± 0.01 | 0.94 ± 0.04 |
| Viperin +TRAF6 | 1.10 ± 0.06 | 1.27 ± 0.01 | 1.15 ± 0.06 |
| Viperin + IRAK1 | 0.70 ± 0.04 | 1.49 ± 0.02 | 2.13 ± 0.13 |
| Viperin +IRAK1+ TRAF6 | 0.40 ± 0.02 | 1.72 ± 0.03 | 4.30 ± 0.22 |
| Viperin +ATP | 1.38 ± 0.07 | 3.48 ± 0.18 | 2.52 ± 0.18 |
| Viperin +GTP | 1.38 ± 0.07 | 1.58 ± 0.11 | 1.14 ± 0.10 |
| Viperin +CTP | 1.38 ± 0.07 | 3.10 ± 0.15 | 2.25 ± 0.16 |
| Viperin +UTP | 1.38 ± 0.07 | 1.57 ± 0.12 | 1.14 ± 0.09 |
| Viperin + IRAK1+ATP | 0.70 ± 0.04 | 4.47 ± 0.11 | 6.39 ± 0.40 |
| Viperin + IRAK1+CTP | 0.70 ± 0.04 | 5.09 ± 0.17 | 7.27 ± 0.48 |
| Viperin +IRAK1+ TRAF6+ATP | 0.40 ± 0.02 | 9.17 ± 1.25 | 22.9 ± 3.3 |
| Viperin +IRAK1+ TRAF6+CTP | 0.40 ± 0.02 | 8.55 ± 0.48 | 21.4 ± 1.6 |

#### Supplementary Figures

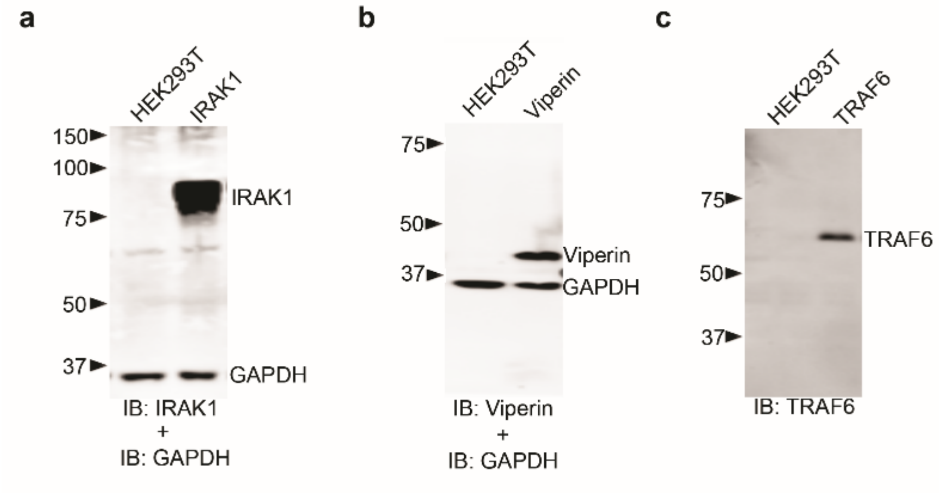


**Supplementary Fig 1.**

**Validation of antibodies used for immuno-blot analysis of viperin, IRAK1 and TRAF6**.

Lysates from non-transfected cells or from transiently transfected cell lines expressing only viperin, IRAK1 or TRAF6 were immunoblotted, probed with antibodies as indicated on the figure panels and subsequently visualized with peroxidase labeled goat anti-rabbit IgG or goat anti-mouse IgG antibodies.

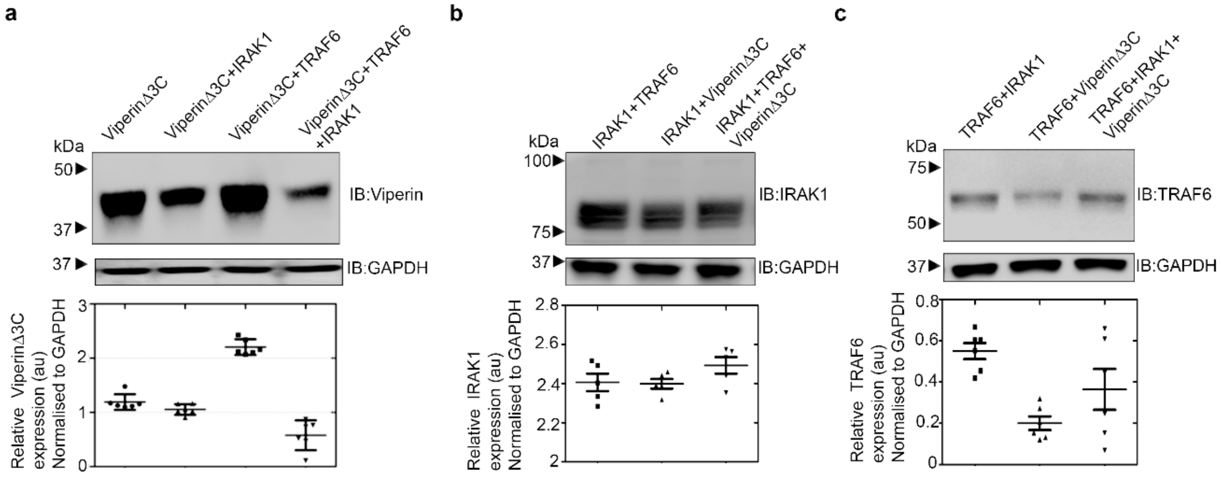


**Supplementary Fig. 2.**

**Effects on relative protein levels of co-expressing catalytically inactive viperin-Δ3C IRAK1 and TRAF6 in HEK293T cells.** HEK293T cells were transiently transfected with indicated expression constructs. 30 h post transfection, cells were harvested and whole-cell lysates from equal numbers of cells were analyzed by immunoblotting using the indicated antibodies. GAPDH served as the loading control.

**a)** Immunoblot analysis for expression of viperinΔ3C in presence and absence of IRAK1 and TRAF6. Co-expression with TRAF6 increases viperin expression whereas co-expression with IRAK1 reduces viperin-Δ3C levels slightly. Co-expression of both TRAF6 and IRAK1 with viperin decreases viperin-Δ3C levels similar to that seen for wildtype viperin.

**b)** IRAK1 levels are not significantly affected by co-expression with viperin-Δ3C or TRAF6

**c)** TRAF6 levels are slightly reduced by co-expression with viperin-Δ3C but not significantly altered by co-expression with IRAK1.

Quantitation of relative protein expression levels for viperin, viperin3C, IRAK1 and TRAF6 relative to GAPDH is presented as mean +/− S.E.M. (n=3) with * indicating $p<0.05$, Student's t-test for independent samples was done.

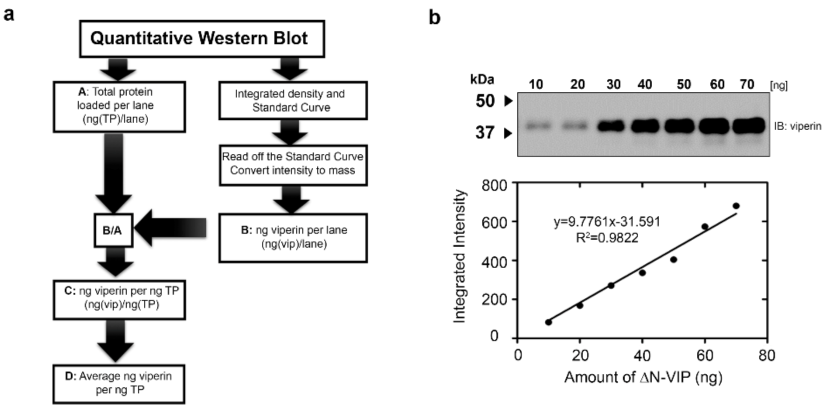


c

| | Viperin (ng) |
|---|---|
| Viperin | 61.8±1.05 |
| Viperin+IRAK1 | 31.07±1.63 |
| Viperin+TRAF6 | 49.28±1.18 |
| Viperin+IRAK1 +TRAF6 | 18.41±0.83 |

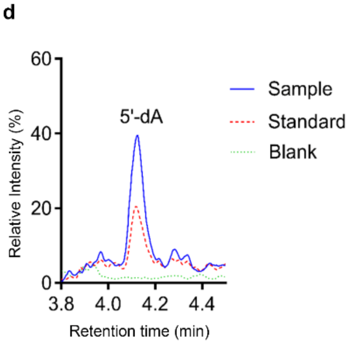


**Supplementary Fig 3.**

**Quantification of viperin in HEK293T cells.**

**a)** Flowchart illustrating the procedure used to determine the amount of viperin in HEK 293T lysates.

**b)** Representative standard curve constructed from increasing concentrations of purified ΔN-viperin, visualize by immunoblotting and quantified .

**c)** Estimation of viperin concentrations in cell lysates using quantitative immunoblot analysis transfected with expression plasmids as indicated. Total protein per cell was calculated by counting cells prior to lysis and by BCA assay. For details see the materials and methods.

**d)** Analysis of 5'dA formed by reductive cleavage of SAM in viperin activity assays. Representative LC-MS chromatograms are shown. For details see supplementary methods.

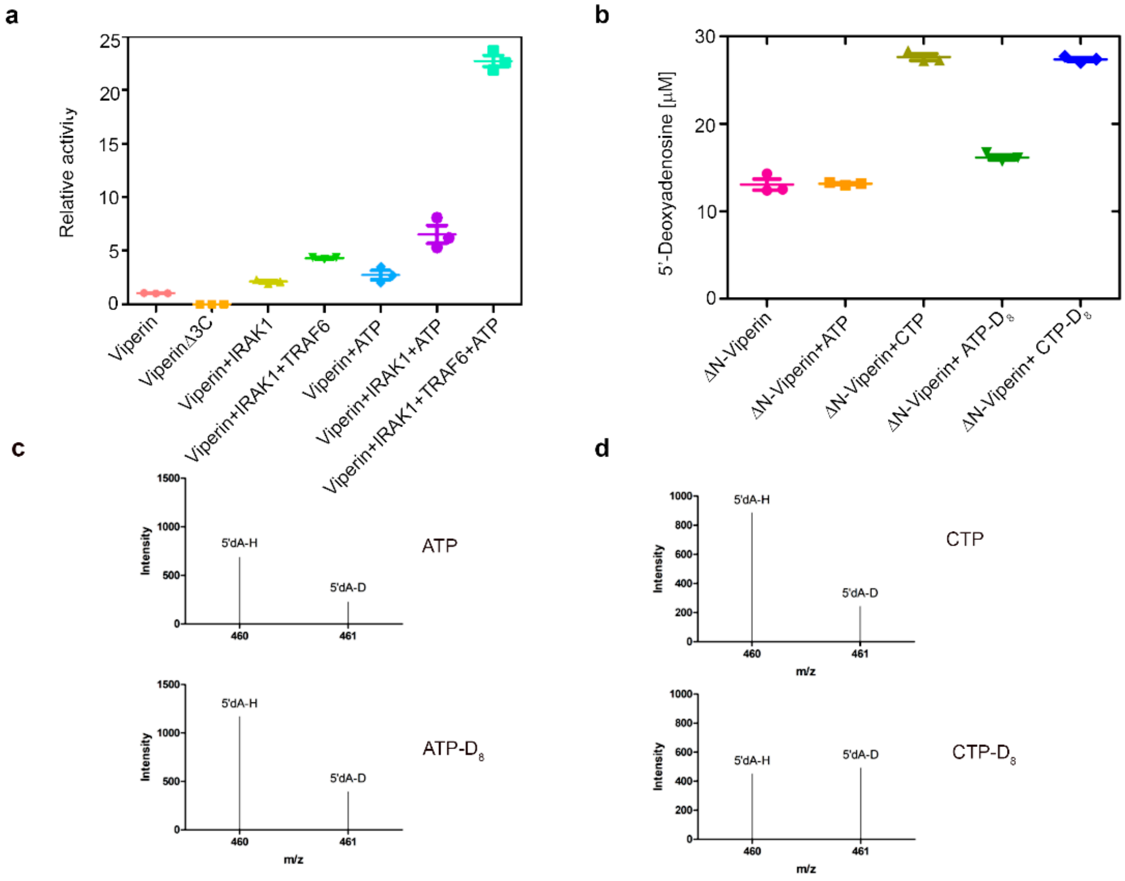


**Supplementary Fig. 4.**

**ATP indirectly stimulates viperin activity in HEK 2983T cell extracts expressing viperin, TRAF6 and IRAK1**

**a)** Viperin activity assayed by detecting the formation of 5′-dA in assays of HEK 293T cell lysates expressing viperin, viperinΔ3C, IRAK1 and/or TRAF6. Relative activity refers to the amount of 5'-dA formed in 1 h, normalized for viperin present in the cell extracts, and relative to the viperin-only sample, which is set to 1.0 on the scale. The data represent the mean and standard deviation of three independent biological replicates with three technical replicates of each measurement. The activity of viperin in cell extracts is stimulated by ATP.

**b)** Comparison of the activity of recombinant ΔN-viperin, purified from *E. coli*, with ATP or with CTP as co-substrates. 5'-dA production was assayed using purified recombinant ΔN-viperin as described in the supplementary methods section. Whereas CTP stimulates the production 5'-dA, ATP does not stimulate viperin activity.

**Supplementary Fig. 4 (cont).**

**c, d)** Viperin catalyzes the abstraction of deuterium from $d_8$-CTP but not ATP**.** HEK293T cell extracts expressing viperin were assayed with ATP, $d_8$-ATP, CTP or $d_8$-CTP and the 5'-dA formed analyzed by MS. 5´-dA formed with $d_8$-CTP as substrate exhibited a characteristic increase in mass indicating the incorporation of deuterium from CTP. 5´-dA formed with $d_8$-ATP as substrate exhibited no increase in mass.

These results indicate that the stimulatory effect of ATP likely derives from the phosphorylation of the endogenous pool of CMP and CDP by phosphoryl transferases in the cell free extracts.

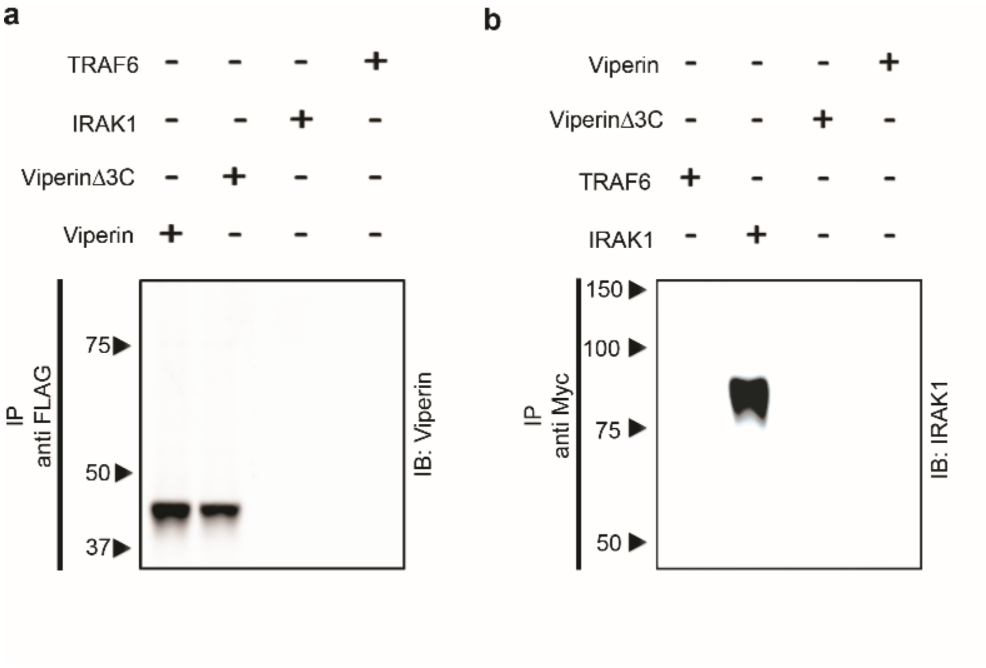


**Supplementary Fig 5.**

**Specificity of antibodies used in pull-down experiments.** No cross reactivity was observed of the anti-FLAG antibodies used to immuno-tag viperin with either IRAK1 or TRAF6. Similarly, no cross reactivity was observed of the anti-Myc antibodies used to immuno-tag IRAK1 with either viperin or TRAF6. Whole cell extracts from HEK 293T were subjected to immunoprecipitation and immunoblotting with the indicated antibodies.

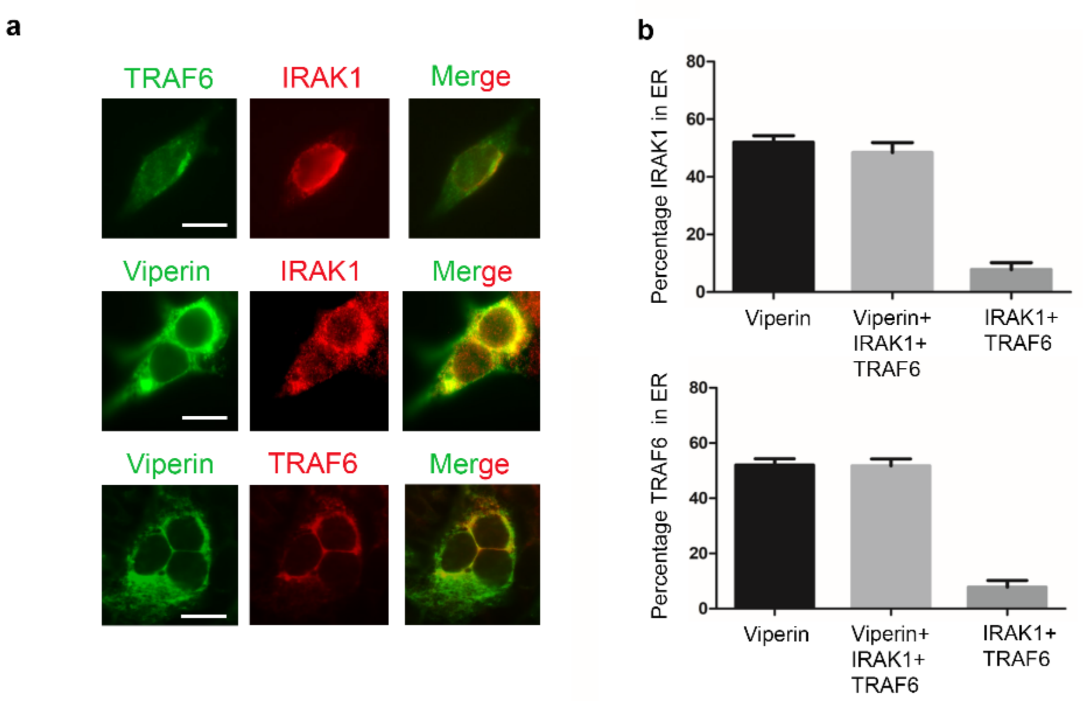


**Supplementary Fig 6.**

**Relocalization of IRAK1 and TRAF6 to the endoplasmic reticulum by viperin.** 293T cells were transiently transfected with viperin and/or IRAK1 and TRAF6. 36 h post transfection, cells were fixed in 4% paraformaldehyde, permeabilized with 0.05% Triton-X, and stained for viperin (red), IRAK1 (red), TRAF6 (red) and calnexin (green) using respective antibodies as described in methods section. Images shown are representative of n = 10 cells. Scale bar = 5 µm.

**a)** *Top panels*: cells co-transfected with IRAK1 and TRAF6 show diffuse expression throughout the cell. *Middle panels*: cells co-transfected with IRAK1, TRAF6 and viperin; staining for viperin (green) and IRAK1 (red) demonstrates co-localization (yellow) of viperin and IRAK1. *Bottom panels*: cells co-transfected with IRAK1, TRAF6 and viperin; staining for viperin (green) and TRAF6 (red) demonstrates co-localization (yellow). Co-expression of IRAK with viperin appears to result in IRAK1 also forming punctate structures that do not co-localize (middle panels). The reason for this is unclear.

**b)** Co-localization was quantified using ImageJ Co-localization plug-in software, presented as a percentage of total viperin/viperinΔ3C/IRAK1/TRAF6 that co-localizes with calnexin. Results are presented as ±SEM of least 10 different cells. ***$P < 0.0001$ (paired $t$ test).

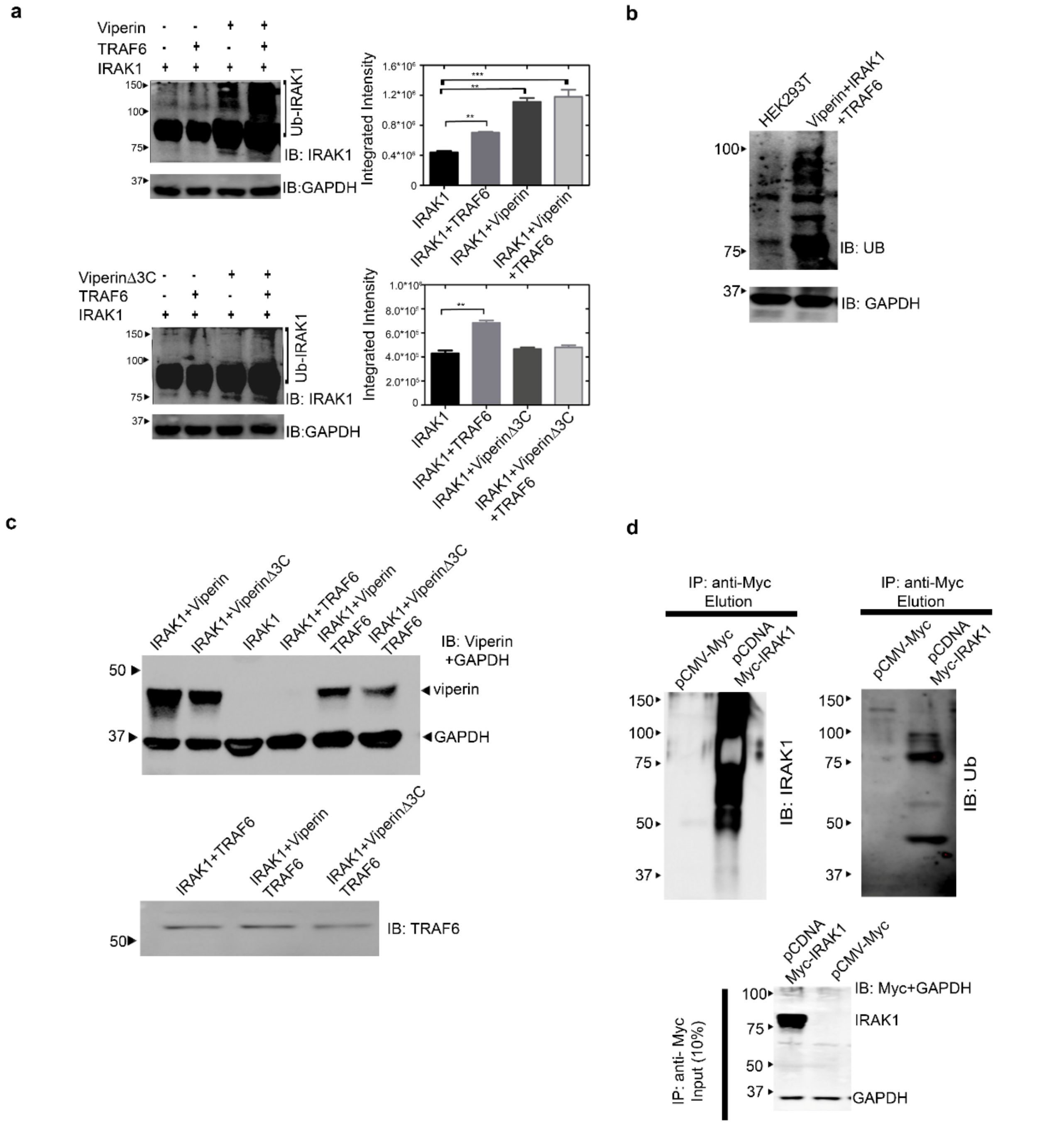


**Supplementary Fig. 7:**

**Control experiments demonstrating that viperin is required for poly-ubiquitination of IRAK1.**

**a)** HEK 293T cells were transfected with the indicated genes and cell lysates subjected to immunoblot analysis using anti-IRAK1 antibody as the probe. GAPDH was used as a loading control. The presence of higher Mr (100 – 150 kDa) immuno-reactive bands indicates poly-ubiquinated forms for IRAK1. The intensity of the IRAK1-reactive bands were calculated for

**Supplementary Fig. 7: (cont)**

each sample by using ImageJ software and compared. *** indicates statistical significance with $p<0.001$.

**b)** Immunoblot analysis of immuno-precipitated IRAK1 from cell lysates of HEK 293T cells transfected with IRAK1, TRAF6 and Viperin which were probed with anti-ubiquitin antibody. The control lane is for non-transfected HEK 293T cells.

**c)** Control immunoblots demonstrating the expression of viperin and TRAF6 in the HEK 293T lysates used in IRAK1 pulldowns. Blots were probed monoclonal antibodies against viperin or TRAF6 using as indicated.

**d)** Control immunoblots demonstrating specificity of Myc-tagged IRAK1 pulldown compared with empty vector control. *Top panels* Immunoprecipitated material analyzed by immunostaining with anti-IRAK1 and anti-ubiquitin antibodies. Bottom Panel Immunostaining of input lysates representing 10% of the amount of IRAK1 used in the pulldown experiments.

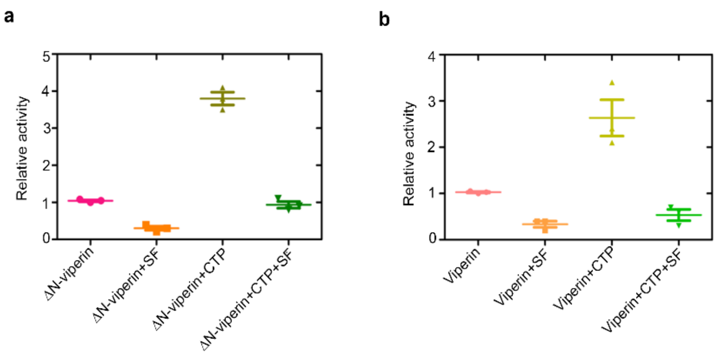


**Supplementary Fig. 8:**

**Sinefungin inhibits viperin activity**.

**a)** Relative activity of purified recombinant ΔN-viperin in the presence of sinefungin (SF), CTP and sinefungin + CTP. The amount of 5'-dA formed in 1 h is plotted relative to the viperin-only sample = 1.0 (uncoupled reductive cleavage of SAM).

**b)** Quantification of viperin activity in transfected HEK 293T cell lysates in the presence of sinefungin (SF), CTP and sinefungin + CTP. The amount of 5'-dA formed in 1 h, normalized for viperin present in the cell extracts, is plotted relative to the viperin-only sample = 1.0.

The data represent the mean and standard deviation of three independent biological replicates with three technical replicates of each measurement. For experimental details see supplementary methods.

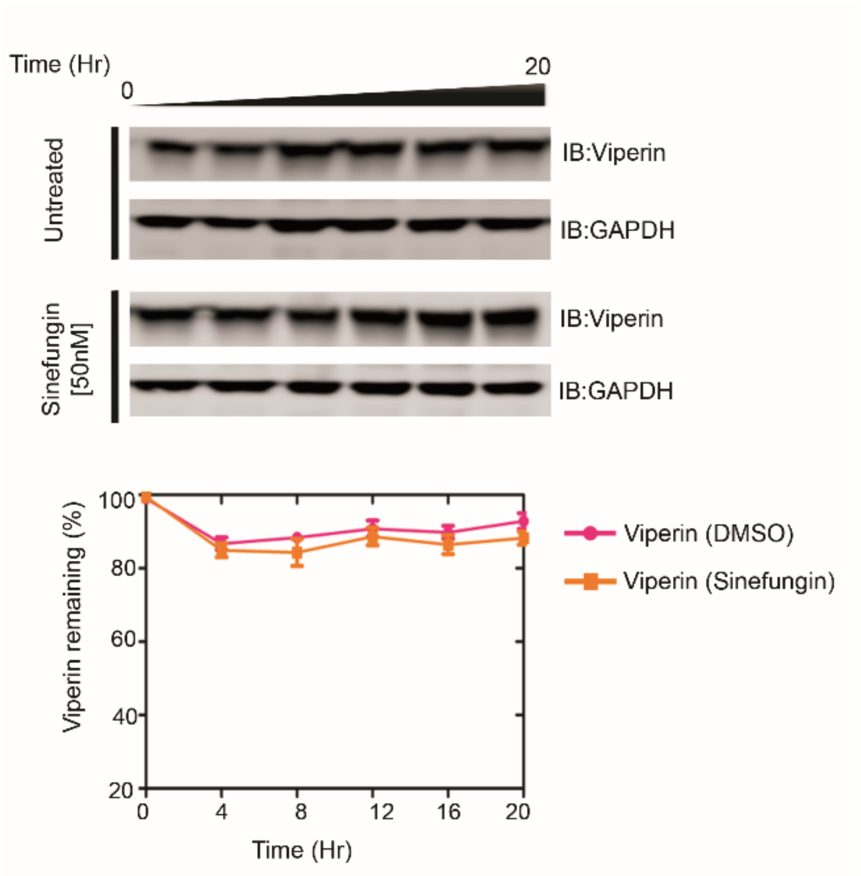


**Supplementary Fig. 9:**

**Sinefungin does not alter wildtype viperin expression levels**. HEK 293T cells were transfected with viperin and were either treated with DMSO or with 50 nM sinefungin 12 h post transfection. At the time points indicated (0, 4, 8, 12, 16, 20 h) cells were harvested and cell lysates were prepared and analyzed by immunoblotting with antibodies as indicated. Band intensities were quantified using a densitometer and plotted as a function of time. Data are expressed as means ± s.d. from three independent experiments. Equal amounts of protein from each sample were immunoblotted with antibody against GAPDH.

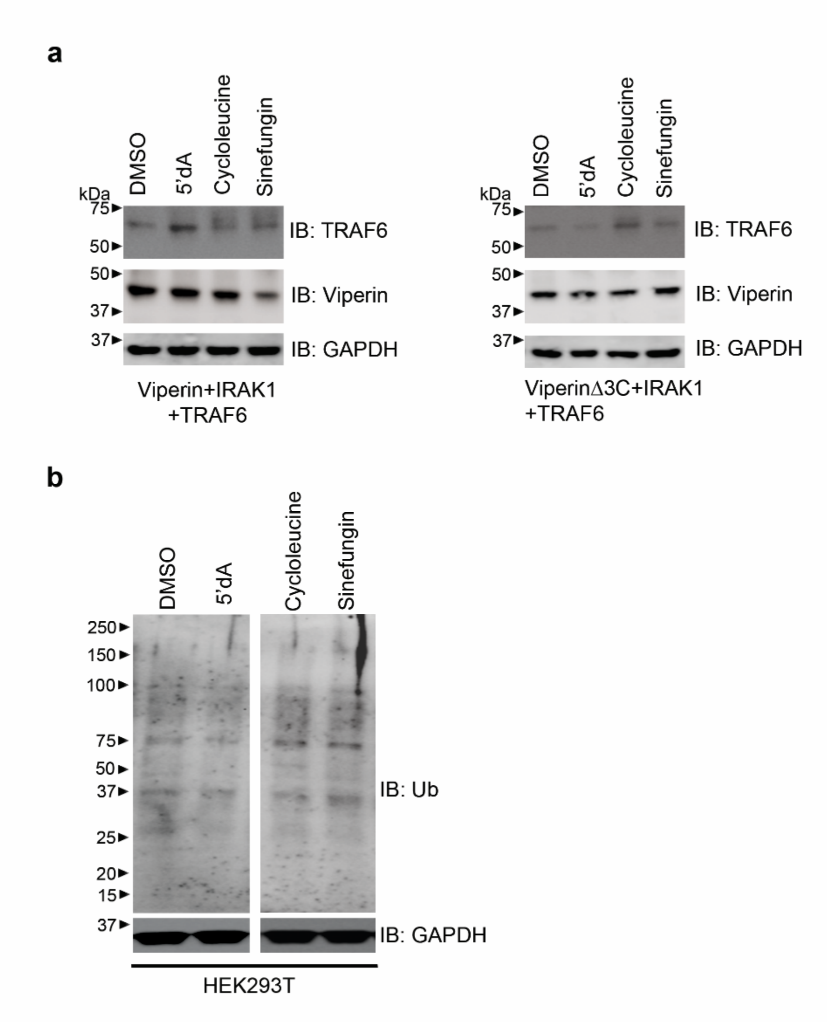


**Supplementary Fig. 10:**

**Analysis of effect of various chemical treatments on cellular expression levels of viperin and TRAF6 and ubiquitination background**.

**a)** The input cell lysates were probed with antibodies as indicated. The different chemical treatments do not significantly change the cellular expression levels for viperin, viperinΔ3C or TRAF6.

**b)** HEK 293T cells were treated with 5'dA, cycloleucine, sinefungin as described in main text and subjected to immunoblotting with anti-ubiquitin antibody. The various chemical treatments do not visibly alter the background of ubiquitinated proteins untransformed HEK 293T cells.
